# A systems approach evaluating the impact of SARS-CoV-2 variant of concern mutations on CD8+ T cell responses

**DOI:** 10.1101/2022.10.21.513200

**Authors:** Paul R. Buckley, Chloe H. Lee, Agne Antanaviciute, Alison Simmons, Hashem Koohy

**Author notes:** These authors contributed equally (Paul R. Buckley, Chloe H. Lee).

## Abstract

T cell recognition of SARS-CoV-2 antigens after vaccination and/or natural infection has played a central role in resolving SARS-CoV-2 infections and generating adaptive immune memory. However, the clinical impact of SARS-CoV-2-specific T cell responses is variable and the mechanisms underlying T cell interaction with target antigens are not fully understood. This is especially true given the virus’ rapid evolution, which leads to new variants with immune escape capacity. In this study, we used the Omicron variant as a model organism and took a systems approach to evaluate the impact of mutations on CD8+ T cell immunogenicity. We computed an ‘immunogenicity potential’ score for each SARS-CoV-2 peptide antigen from the ancestral strain and Omicron, capturing both antigen presentation and T cell recognition probabilities. By comparing ancestral vs. Omicron immunogenicity scores, we reveal a divergent and heterogeneous landscape of impact for CD8+ T cell recognition of mutated targets in Omicron variants. While T cell recognition of Omicron peptides is broadly preserved, we observed mutated peptides with deteriorated immunogenicity that may assist breakthrough infection in some individuals. We then combined our scoring scheme with an *in-silico* mutagenesis, to characterise the position- and residue-specific theoretical mutational impact on immunogenicity. While we predict many escape trajectories from the theoretical landscape of substitutions, our study suggests that Omicron mutations in T cell epitopes did not develop under cell-mediated pressure. Our study provides a generalisable platform for fostering a deeper understanding of existing and novel variant impact on antigen-specific vaccine- and/or infection-induced T cell immunity.

## Introduction

Cellular and humoral responses are pillars of adaptive immunity following vaccination or natural SARS-CoV-2 infection. A clear understanding of how emergent variants of concern (VOC) affect adaptive immunity is essential for effective control of the pandemic, especially for a virus with such dynamic evolution, which threatens a future VOC with additional escape capacity.

Upon its emergence, the SARS-CoV-2 VOC Omicron BA1 caused global concern due to alarming mutations in its surface glycoprotein. It however became clear that Omicron and derivative subvariants BA2, 4, 5 among others are broadly associated with milder disease outcomes than previous strains [1]. Nevertheless, their higher transmissibility rates[2] contributed to widespread infection, social disruption and healthcare turbulence.

While numerous studies have illustrated extensive humoral immune escape by Omicron and its subvariants[3–6], accumulating reports indicate that broadly, mutations in Omicron and other current VOCs do not drastically hinder T cell responses[7,8]; likely due to the diversity of HLAs and the breadth of epitopes targeted by T cells. To this end, a series of studies focused on memory T cells have shown that impact of variant mutations on T cell response is limited and the majority of CD4+ and CD8+ T cell responses are preserved in most naturally infected and/or vaccinated individuals[9–15]. For Omicron specifically, Tarke et al reported preservation of at least 84% and 85% for CD4+ and CD8+ cell response respectively following a range of different vaccines, whereas a highly significant decrease for memory B cells was observed[16]. These studies have offered great insights that have informed policymaking on mitigation of the disease e.g., booster rollouts.

Nevertheless, multiple studies have demonstrated substantial impairment in some cases [17,18]. Indeed, Naranbhai et al found that in ∼20% of individuals (n=10) examined, >50% of their T cell responses were impaired after Omicron [17]. More recently, Reynolds et al. found that individuals infected in the first wave and subsequently with Omicron had particularly poor T cell responses[18] and that Omicron BA1 was a poor booster of immunity to subvariant Omicron infections. Furthermore, Suryawanshi et al. found that infection and vaccination ‘hybrid’ immunity does not appear to protect against different variants [19]. Indeed, such data are beginning to demonstrate complex heterogeneity underpinning whether breakthrough infections occur; and that the extent to which mutations from VOC can impact T cell immunity in individuals is varied.

Adding to this heterogeneous landscape, is that although many individuals now experience COVID-19 with mild symptoms, there are increasing reports of complications following SARS-CoV-2 infection. For example, considerable numbers of individuals suffer with post- acute sequalae of COVID-19[20–22] (long COVID) and there are increasing reports of multisystem inflammatory disorders and severe acute hepatitis, where SARS-CoV-2 has been implicated as the causative agent [20,23–25]. Indeed, while T cells are thought to provide a barrier against Omicron infection where antibody responses are evaded, there is increasing evidence that T cells may on the other hand contribute to post-COVID complications in some individuals, e.g., through provoking inflammatory disorders[20,23,26,27], elevated T cell exhaustion in long COVID[21] and CD8+ T cell infection and lymphopenia following SARS-CoV-2 exposure [28].

Collectively, these studies raise diverse questions regarding heterogeneity underpinning how T cells may protect from or exacerbate COVID-19 and related diseases. One such question, is how mutations in VOCs such as Omicron have impacted the landscape of SARS-CoV-2- specific T cell responses. Additionally, the high prevalence of humoral and cellular immunity from vaccination and/or natural infection may impose selection pressures promoting new variants, threatening a future strain characterised by T cell escape. Taken together, these studies demonstrate it is vital to understand how mutations from existing VOCs such as Omicron – and theoretical mutations – affect T cell responses to SARS-CoV-2.

Multiple studies have begun to address this gap [8,29–31]. For example, Nersisyen et al [29] recently compared all theoretical HLA ligands from the Wuhan Hu-1, Delta and Omicron proteomes. The authors concluded that most HLA alleles and theoretical MHC haplotypes are not significantly affected by mutations in Delta / Omicron; therefore, effective T cell immunity is likely to be maintained. Notwithstanding, the authors reported a reduction in ‘tightly binding’ spike-derived HLA-B*07:02 ligands for both Delta and Omicron and an elimination of ligands for HLA-DRB1-03:01. Indeed, studies to date have generally focused on comparing antigen presentation of HLA ligands across VOCs. However, while antigen presentation is *required* for T cell immunogenicity, it is not *sufficient*. Therefore, to foster a detailed understanding of how mutations can affect SARS-CoV-2-specific T cell responses, it is vital to examine the subset of MHC-bound ligands that are known to invoke T cell activation.

Dolton et al., recently argued that the range of potential mutational impact on SARS-CoV-2- specific T cell recognition is poorly understood and urgently requires further study[32]. Indeed, despite the great insights from the above research, a systematic study to examine the impact of mutations at all known SARS-CoV-2-specific CD8+ T cell targets, on T cell immunogenicity has not yet been reported. Therefore, there is an unmet need for research to evaluate and infer the impact of current and theoretical future mutations on T (and B) cell immunity.

In this study, using SARS-CoV-2 CD8+ T cell epitopes as a model system, we present a systematic framework to evaluate the impact of mutations at T cell targets on their immunogenicity (Fig 1A). To model peptide immunogenicity, we consider i) antigen presentation predictions using netMHCpan, and ii) T cell recognition by leveraging a deep neural network workflow we have recently developed for accurate predictions in this setting (Fig. 1A). By inferring the effects of Omicron mutations on CD8+ T cell epitopes, we found a heterogeneous landscape of impact, encompassing the most likely targets for T cell escape, as well as pMHC with potentiated immunogenicity following mutation. We then present an *in-silico* mutagenesis and dissect the impact of theoretical substitutions on SARS-CoV-2-specific CD8+ T cell responses. Our approach allows systematic evaluation of the impact of mutation on T cell responses, not only for those conferred by existing variants such as Omicron, but also theoretical mutations. We believe that such an approach will evolve into pipelines to predict the impact of emerging variants on T cell immunity, e.g., by estimating the probability of T cell immune escape. These concepts could additionally be applied to other diseases that are affected by pathogenic mutations.

**Figure 1:**
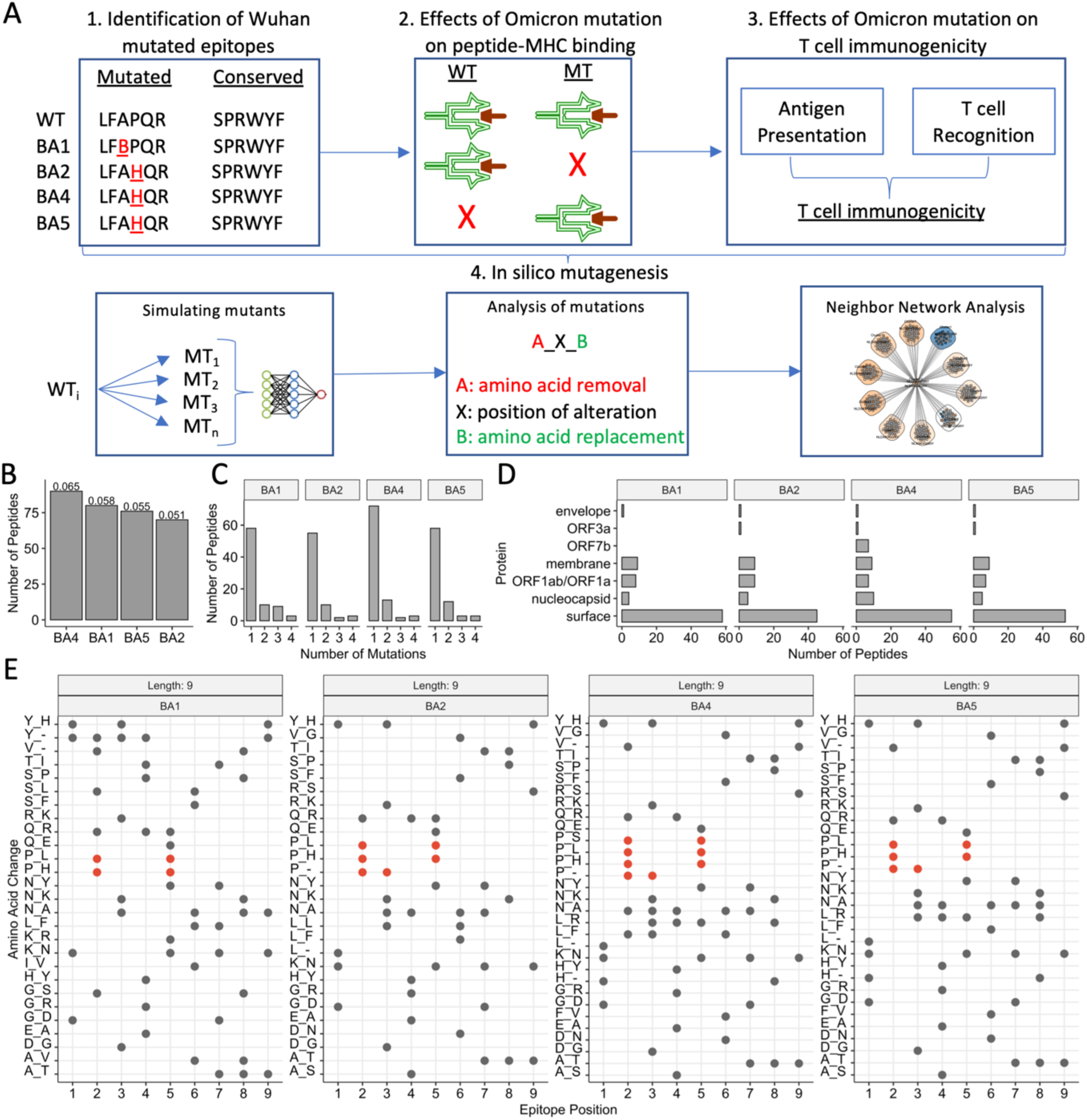
Study Overview and Identification of SARS-CoV-2-specific CD8+ T cell targets with mutation(s) in Omicron and its subvariants. A) Overview of the Study: 1) First, we identified the set of ‘Wuhan-mutated’ epitopes: CD8+ T cell targets from Wuhan with mutations in Omicron (BA1) or its subvariants (BA2, BA4, BA5). 2) Using netMHCpan 4.1, we analysed the effects of mutations from aforementioned variants of concern on antigen presentation, through comparing predicted binding metrics (affinity *nM*, normalised binding affinity rank score) between Wuhan variant CD8+ T cell epitopes and counterpart mutant epitopes. To appropriately analyse potential bi-directional changes in antigen presentation between wildtype and mutant peptides, we incorporated wildtype-mutant (WT-MT) paired samples where top) both WT and MT bind MHC *k*, middle) where WT but not MT binds MHC *k*, bottom) where WT was not a ligand for MHC *k* in Wuhan-Hu-1 but the MT now binds MHC. 3) Next, we analysed the effects of existing variant of concern mutations on T cell immunogenicity. T cell immunogenicity incorporates two key components: antigen presentation and T cell recognition. We therefore combined two scores for this analysis: i) antigen presentation – generated by netMHCpan 4.1 – and ii) T cell recognition, generated by an in-house deep learning model ‘TRAP’. 4) We next performed a comprehensive *in silico* mutagenesis to examine the effects of theoretical mutations on T cell immunogenicity. To do this, we first simulated each mutant (MT_1_‥MT_n_) that can be generated via single point substitutions for each WT Wuhan-mutated peptide of interest (WT_i_). For each wildtype and mutant peptide, we generated an immunogenicity score. Next, we analysed the effects of mutation on immunogenicity, through analysing WT-MT paired changes in immunogenicity score. In this paper, we follow nomenclature where a ‘full’ mutation comprises three components: ‘A_x_B’. Here, residue ‘A’ is removed from epitope sequence position *x* and is replaced with residue ‘B’. Finally, we adopted the ‘Neighbor Network’ framework of Ogishi et al. to visualise the effects of different residue substitutions, to help identify the most escape-prone mutations and understand how different mutations in different positions affect each epitope. B) The number of CD8+ T cell targets which exhibit at least one mutation from BA1 Omicron and/or derivative subvariants BA2, BA4, BA5. Numeric labels show the frequency of mutated CD8+ T cell epitopes, given a particular variant compared with the total number of assessed CD8+ T cell targets (n=1380). C) The distribution of the number of mutations found in CD8+ T cell targets across assessed variants of concern (with respect to Wuhan Hu-1). D) The distribution of originating proteins for SARS-CoV-2 Wuhan Hu-1 CD8+ T cell targets with a mutation in each variant. E) The landscape of amino acid substitutions across different variants of concern within CD8+ T cell targets of length 9 residues. ‘X_Y’ on the y-axis indicates the removal of amino acid X, which is replaced by amino acid Y. The x-axis shows the position in the epitope that the mutation is observed. Red points highlight amino acid alterations which remove a ‘Proline’.

## Results

### The subset of Wuhan Hu-1 CD8+ T cell epitopes with mutation in Omicron

To investigate the effect of existing mutations on SARS-CoV-2-specific CD8+ T cell epitopes, we curated a pool of 1380 unique SARS-CoV-2 peptides from epitope databases that have been functionally evaluated for CD8+ T cell response (see Methods). Of these, 9-mers were most common (∼54%), followed by 10-mers (25.6%) (S1A).

To identify immunogenic Wuhan Hu-1 epitopes which are mutated in the parent Omicron variant BA1 and subvariants BA2, BA4 and BA5, we mapped our pool of 1380 peptides to each respective proteome (see Methods). We observed that BA4 produced the greatest number of mutations in unique Wuhan Hu-1 CD8+ T cell targets (90/1380, ∼6.5%), followed by BA1 (80/1380, ∼5.8%), BA5 (76/1380, ∼5.5%) and BA2 (70/1380, ∼5%) (Fig. 1B). For each variant, most of these alterations were single point mutations, although epitopes with 2 or 3 mutations were also relatively abundant (Fig 1C, Table 1). Unsurprisingly, for each variant, most CD8+ T cell epitopes that exhibited mutations were derived from spike glycoprotein (Fig. 1D).

**Table 1:**
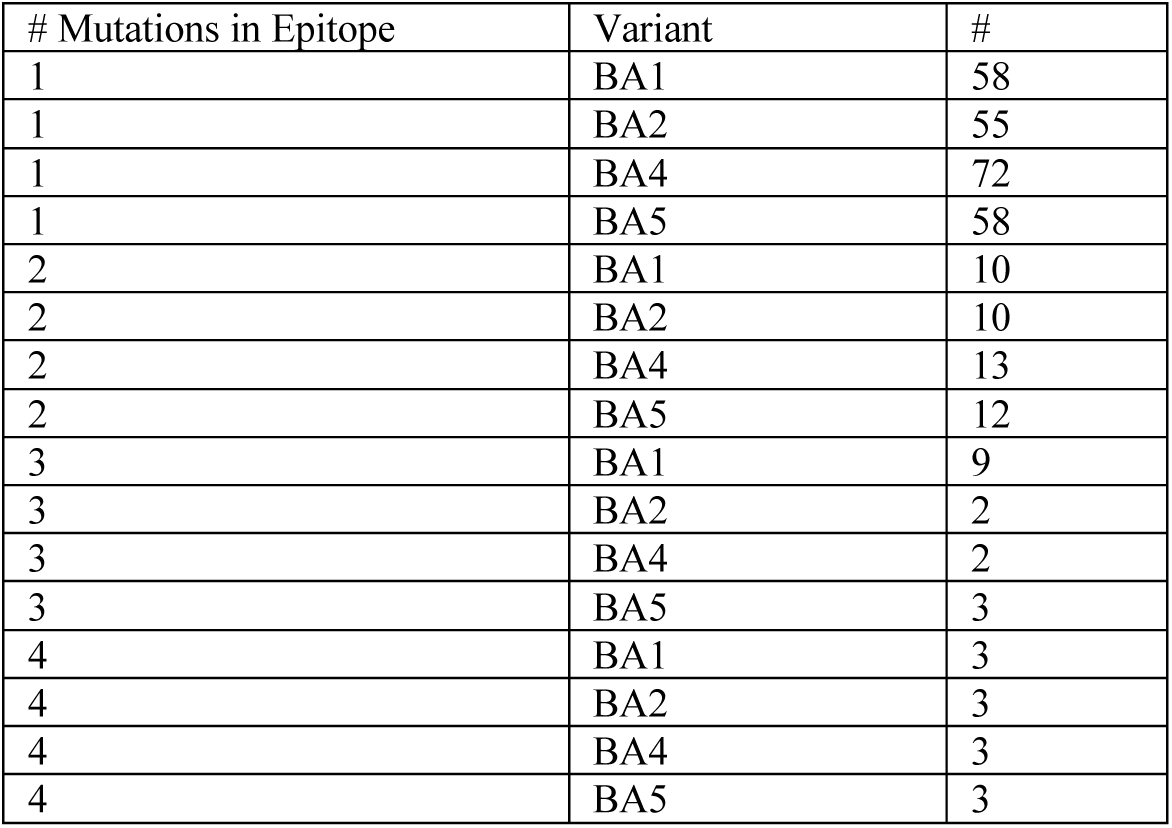
The number of epitopes (#) with *x* number of mutations in CD8+ T cell regions of variants BA1, BA2, BA4, BA5, with respect to Wuhan Hu-1.

| # Mutations in Epitope | Variant | # |
| --- | --- | --- |
| 1 | BA1 | 58 |
| 1 | BA2 | 55 |
| 1 | BA4 | 72 |
| 1 | BA5 | 58 |
| 2 | BA1 | 10 |
| 2 | BA2 | 10 |
| 2 | BA4 | 13 |
| 2 | BA5 | 12 |
| 3 | BA1 | 9 |
| 3 | BA2 | 2 |
| 3 | BA4 | 2 |
| 3 | BA5 | 3 |
| 4 | BA1 | 3 |
| 4 | BA2 | 3 |
| 4 | BA4 | 3 |
| 4 | BA5 | 3 |

Leveraging our dataset of Wuhan Hu-1 immunogenic epitopes with a mutation in Omicron strains (hereby referred to as ‘Wuhan-mutated’) and their respective variant counterparts, we explored alterations at different sequence positions across 9-mer (Fig. 1E) and 10-mer (S1B) peptides. We observed P→L/H mutations at P2 of 9-mers among all variants. BA4 poses an additional P→S mutation in this position. For 10-mers, we also observed proline substitutions, although they were less common (S1B).

Proline substitutions are of particular interest, as a P→L mutation (P4, YLQ<u>P</u>RTFLL-HLA- A02: in non-Omicron variants) has recently been shown to escape SARS-CoV-2-specific CD8+ T cell responses[32]. Furthermore, a report by Hamelin et al. found that proline substitutions arise from mutational bias in SARS-CoV-2 evolution, which compromised binding of HLA-B07 ligands[33]. The underlying nucleotide mutation (C→U) has been shown to possess a close relationship with HIV-1 T cell immunity. Taken together, mutations in globally disseminated VOCs are consistent with alterations that are known to compromise T cell epitopes.

### Effects of Omicron mutations on peptide-MHC-I binding

While antigen presentation by MHC is not sufficient for an effective T cell response, it is a pre- requisite for T cell recognition. We therefore used the known CD8+ T cell targets in Wuhan to systematically study the impact of existing mutations on their presentation status. In this section, focusing on immunogenic Wuhan Hu-1 CD8+ T cell epitopes with alterations in Omicron variants, we report the effects of mutations on their capacity to bind MHC.

We first predicted HLA binding metrics (see Methods) of each Wuhan-mutated peptide to 64 HLAs which i) are commonly observed in epitope datasets and ii) have previously been employed by the ‘TCoV’ pipeline to compare overall MHC binding of SARS-CoV-2 variants [29]. As we have shown, such a strategy can identify MHC ligands with 98% accuracy [2]. Here, HLA-A*29:02 is predicted to present the most Wuhan Hu-1 CD8+ T cell targets (309/1380), while HLA-B*27:05 is predicted to present the least (35/1380).

Expanding this approach, we next compared the antigen presentation status of Wuhan-mutated and their BA1, BA2, BA4, or BA5 Omicron counterpart peptides against these 64 HLA-I alleles (see Methods). By comparing 463 spike-derived, paired WT vs. MT predicted pMHC (57 peptides), we observed that BA1 Omicron mutants were predicted as weaker binders (higher nM) to MHC-I alleles than their Wuhan Hu-1 counterparts (Fig 2A). Different HLAs bind their ligands in different nM ranges, thus by analysing *netMHCpan* rank scores we confirmed these results are not due to HLA biases in the dataset (S2A). We found a similar albeit weaker trend after comparing CD8+ T cell targets across *all* SARS-CoV-2 proteins (S2B). We also observed that BA2, BA4 and BA5 mutants also exhibited weaker predicted binding to MHC-I compared with Wuhan Hu-1 (Fig 2B and S2C-D).

**Figure 2:**
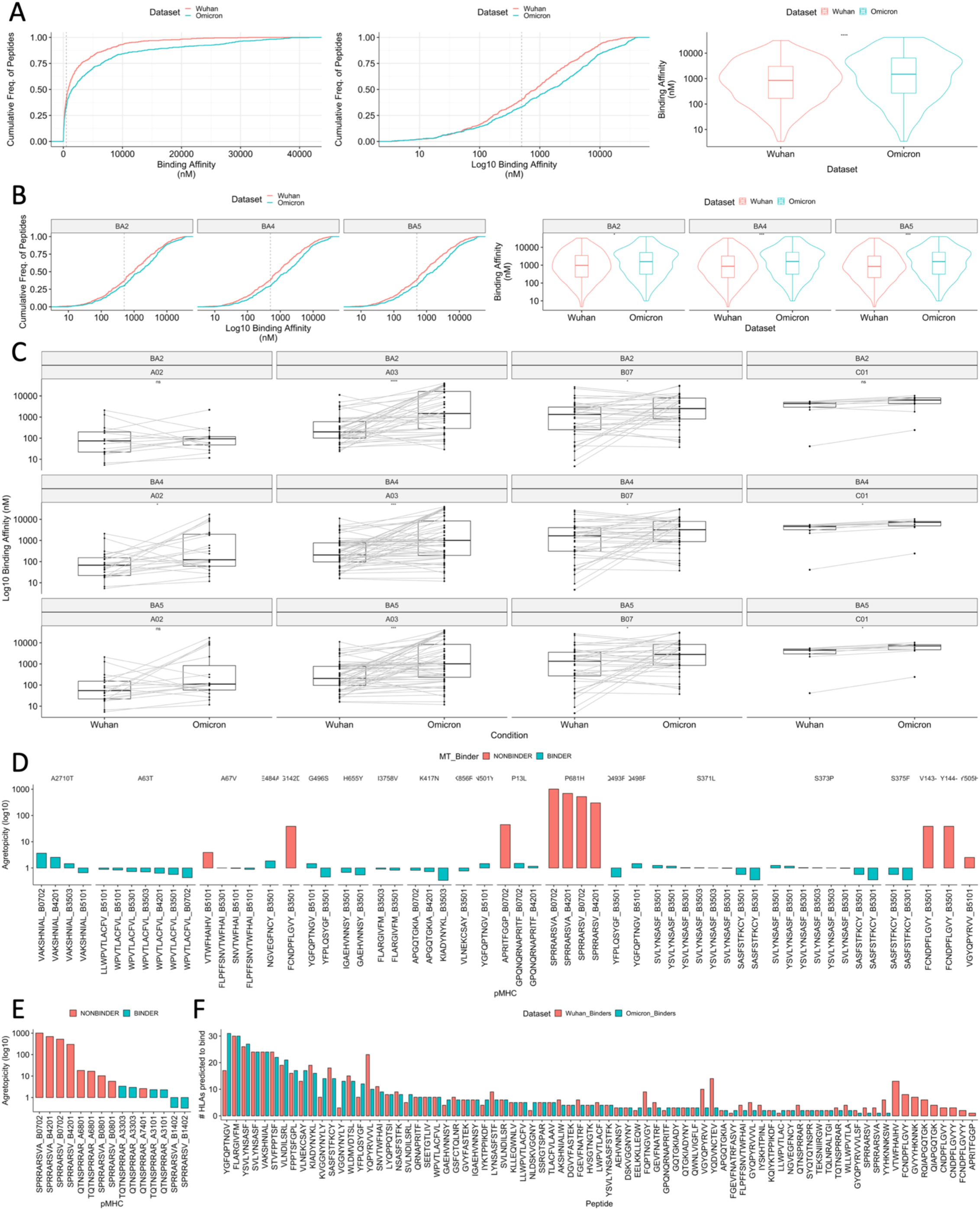
The impact of BA1 Omicron and subvariant mutations on HLA binding of mutated SARS-CoV-2 CD8+ T cell targets. HLA binding predictions were made using netMHCpan 4.1. We analysed any paired wildtype (WT) or mutant (MT) samples where *either* the WT or the MT were predicted to bind MHC, to incorporate retained binding status and bi-directional transitions between WT and MT. Analyses were performed for the set of Wuhan Hu-1 CD8+ T cell targets with a mutation and their mutated counterparts. A) empirical cumulative distribution (ecdf) plot showing i) the cumulative frequency of peptides given the range of binding affinities (nM) for Wuhan-mutated vs BA1 Omicron pMHC, ii) the cumulative frequency of peptides, given the range of log10 transformed binding affinities (nM) for Wuhan-mutated vs BA1 Omicron pMHC. iii) violin plots comparing the distribution of binding affinities (nM) between Wuhan-mutated pMHC vs. BA1 Omicron mutants. Significance was assessed using a Wilcoxon rank test. Dashed line shows 500nM. B) left) log10 transformed ecdf plots and right) violin plots comparing the binding affinity (nM) of Wuhan-mutated vs mutant pMHC, where the mutated epitope is derived from variants BA2, BA4, BA5. C) Paired boxplots for different HLA supertypes (HLA-A02, HLA-A03, HLA-B07, HLA-C01), comparing the binding affinities (nM, log 10) between Wuhan-mutated and mutant pMHC where the mutated epitope is derived from variants BA2, BA4, BA5. Significance was assessed using paired Wilcoxon rank tests. D) Barplots showing log 10 scaled agretopicities (MT nM/WT nM), for BA1 Omicron mutations affecting HLA-B07 supertype pMHC. Agretopicity>0 indicates a detriment in HLA binding for the mutant compared with the wild type. Plots are color labelled by whether the mutant pMHC is predicted to bind MHC or not (red). Red bars therefore indicate those mutant CD8+ T cell pMHC predicted to no longer bind MHC following variant mutation. E) Barplots showing log10 scaled agretopicities for BA1 mutations within CD8+ T cell targets that contain the spike glycoprotein motif ‘PRRA’. ‘PRRA’ is a motif unique to SARS-CoV-2, which is hypothesised to form a portion of a putative core of a SARS-CoV-2 superantigen. Plots are color labelled by whether the mutant pMHC is predicted to bind MHC or not (red). F) A barplot showing the number of HLAs predicted to bind each mutated wildtype epitope (labelled: Wuhan_Binders) vs. the Omicron BA1 counterpart (labelled: Omicron_Binders).

By grouping paired observations by HLA supertype, we observed that binding affinity for BA1 spike-derived B07-pMHC is perhaps weaker compared to Wuhan Hu-1 pMHC, although this effect was not significant (S2E). For BA2 and BA5, we found that ligands bound to HLA-A03 and HLA-B07 were significantly impaired (Fig 2C), with the addition of HLA-A02 for BA4. As ∼25-35% of the global population possess an A02, A03 or B07 supertype allele[33,34], it is plausible that binding detriments for particular pMHC may lead to impairments in T cell reactivity for individuals carrying certain HLA, with differences across subvariants. Our data are consistent with Hamelin’s arguments that mutational biases may shape SARS-CoV-2 T cell reactivity, and we show that mutations from globally spread VOC in B07-binding CD8+ T cell target regions are predicted to significantly weaken pMHC binding affinity.

Focusing on B07-pMHC with mutations in the original BA1 strain, we observed that the overlapping ligands ‘SPRRARSVA’ and ‘SPRRARSV’ are most detrimentally affected (Fig 2D). These CD8+ T cell targets are impacted by a P→H mutation in the P2 anchor position, consistent with Hamelin’s work. Interestingly, the Wuhan Hu-1 sequences of these epitopes contain ‘PRRA’, which is a key motif forming a putative core of a hypothesised SARS-CoV- 2 ‘superantigen’ with structural similarities to the prototypical superantigen - staphylococcal enterotoxin B (SEB)[26,27]. This SARS-CoV-2 superantigen is hypothesised to play a role in post-COVID-19 multisystem inflammatory syndrome and the recent rise in children with severe acute hepatitis[20,23]. In fact, 9/15 (60%) pMHC-I containing this hypothesised superantigen core are predicted to become ‘nonbinders’ as a result of BA1 mutation (Fig 2E, Table 2). Given the global spread of Omicron and its subvariants, these data suggest that CD8+ T cell responses induced by this hypothesised superantigen may currently be limited to few HLA (HLA-A*33:03, −A31:03, −B14:02) which remain as presenters following Omicron mutation.

**Table 2:**
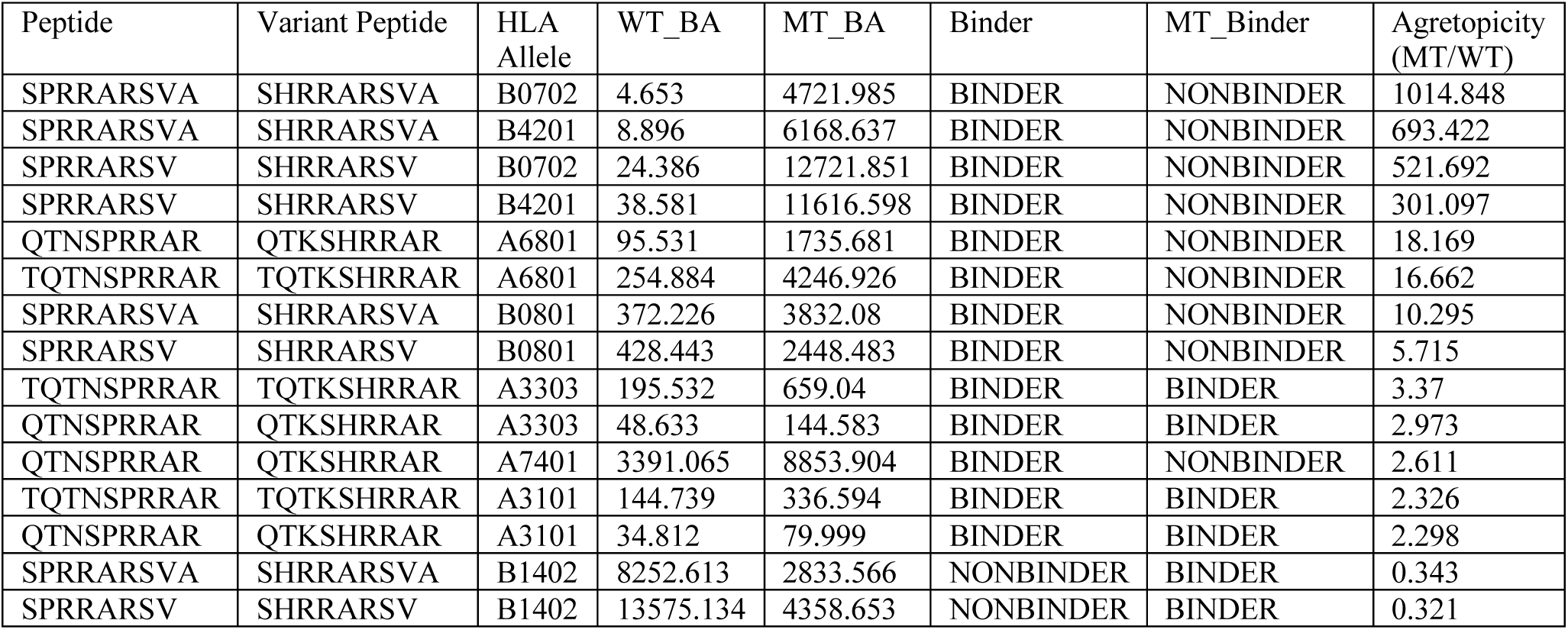
CD8+ T cell targets containing the motif ‘PRRA’ with a mutation in Omicron BA1 (with respect to Wuhan Hu-1). ‘PRRAR’ motif is hypothesised to be a key insert of a core of a SARS-CoV-2 ‘superantigen’. ‘Peptide’ lists the wildtype Wuhan Hu-1 CD8+ T cell target, ‘Variant Peptide’ lists the corresponding identified mutant in BA1, ‘HLA Allele’ lists for each Peptide-Variant, HLAs where either the wildtype peptide or the mutant (Variant Peptide) is predicted to bind, ‘WT_BA’ and ‘MT_BA’ shows the predicted binding affinity of the wildtype and mutant peptides respectively. ‘Binder’ and MT_Binder’ show whether the wildtype or mutant is predicted to bind the listed HLA (determined by binding affinity rank threshold output by netMHCpan 4.1). ‘Agretopicity’ shows the differential agretopicity index, the ratio (MT BA/WT BA) of mutant to wildtype binding affinities. An agretopicity >0 indicates an impairment in binding affinity for the mutant compared with the wildtype while agretopicity <0 indicates the wildtype binds with lower affinity.

| Peptide | Variant Peptide | HLA Allele | WT_BA | MT_BA | Binder | MT_Binder | Agretopicity (MT/WT) |
| --- | --- | --- | --- | --- | --- | --- | --- |
| SPRRARSVA | SHRRARSVA | B0702 | 4.653 | 4721.985 | BINDER | NONBINDER | 1014.848 |
| SPRRARSVA | SHRRARSVA | B4201 | 8.896 | 6168.637 | BINDER | NONBINDER | 693.422 |
| SPRRARSV | SHRRARSV | B0702 | 24.386 | 12721.851 | BINDER | NONBINDER | 521.692 |
| SPRRARSV | SHRRARSV | B4201 | 38.581 | 11616.598 | BINDER | NONBINDER | 301.097 |
| QTNSPRRAR | QTKSHRRAR | A6801 | 95.531 | 1735.681 | BINDER | NONBINDER | 18.169 |
| TQTNSPRRAR | TQTKSHRRAR | A6801 | 254.884 | 4246.926 | BINDER | NONBINDER | 16.662 |
| SPRRARSVA | SHRRARSVA | B0801 | 372.226 | 3832.08 | BINDER | NONBINDER | 10.295 |
| SPRRARSV | SHRRARSV | B0801 | 428.443 | 2448.483 | BINDER | NONBINDER | 5.715 |
| TQTNSPRRAR | TQTKSHRRAR | A3303 | 195.532 | 659.04 | BINDER | BINDER | 3.37 |
| QTNSPRRAR | QTKSHRRAR | A3303 | 48.633 | 144.583 | BINDER | BINDER | 2.973 |
| QTNSPRRAR | QTKSHRRAR | A7401 | 3391.065 | 8853.904 | BINDER | NONBINDER | 2.611 |
| TQTNSPRRAR | TQTKSHRRAR | A3101 | 144.739 | 336.594 | BINDER | BINDER | 2.326 |
| QTNSPRRAR | QTKSHRRAR | A3101 | 34.812 | 79.999 | BINDER | BINDER | 2.298 |
| SPRRARSVA | SHRRARSVA | B1402 | 8252.613 | 2833.566 | NONBINDER | BINDER | 0.343 |
| SPRRARSV | SHRRARSV | B1402 | 13575.134 | 4358.653 | NONBINDER | BINDER | 0.321 |

By analysing paired changes in binding affinity for individual Wuhan-mutated peptides, we found substantial variation (S2F), where some peptides, e.g., FQPT*, FCND*, YQP*, experienced considerable detriments to their binding capacity, while some are unchanged or perhaps exhibited stronger affinity (e.g., KVG*).

Our observations thus far suggest impairments in individual T cell reactivity against VOCs as observed by Naranbhai et al. may be conditioned by HLA genotype and bias toward certain epitopes amongst an individual’s responding T cell compartment. Indeed, by analysing the number of HLAs predicted to present each peptide before (wild type, red) and after (mutant, blue) Omicron mutation, we found that these alterations are predicted to remove 9 epitopes (∼11% of mutated CD8+ T cell targets) as predicted HLA class I ligands entirely (Fig 2F). Although this represents a small proportion (9/1380) of the total pool of immunogenic SARS-CoV-2 CD8+ T cell epitopes, it is plausible that individuals with memory responses biased toward such epitopes may have impaired T cell responses to Omicron and possibly breakthrough infections.

Overall, we have observed that VOC mutations have the capacity to disrupt HLA binding to functionally evaluated SARS-CoV-2-specific CD8+ T cell targets, which, for epitopes containing a hypothesized SARS-CoV-2 superantigen motif, has led to a narrower MHC-binding repertoire. Our data suggests that the landscape of presenting HLAs is also impacted by the mutations. While Omicron mutations are only observed in small proportions of total Wuhan Hu-1 CD8+ T cell targets, given these collective data, it is plausible that patients with certain HLA and/or memory responses biased toward certain pMHC, could be more affected by SARS-CoV-2 VOC mutations than the current paradigm suggests.

### Comparison of Wuhan and Omicron T cell immunogenicity

Peptide presentation by MHC molecules, although necessary, is insufficient to infer T cell immunogenicity in humans[35]. To invoke a T cell response, a presented pMHC must be recognised by a cognate T cell. Therefore, to build on previous insights into HLA binding, we examined how mutations in Omicron VOCs can affect the immunogenicity potential (i.e., both HLA binding and T cell recognition) of pMHC. To do this, we combined two prediction scores for each peptide-MHC: 1) antigen presentation potential and 2) T cell recognition potential (see Methods).

While *in silico* models that predict antigen presentation can be highly accurate (such as netMHCpan), state-of-the-art models predicting the subset of HLA ligands that then invoke T cell responses possess limited accuracy[36]. We recently developed TRAP, a Convolutional Neural Network (CNN) model that offers improved predictions of T cell recognition potential of HLA-I presented 9- and 10-mer peptides. To predict the T cell recognition potential of Wuhan vs. Omicron pMHC of interest, we trained an instantiation of TRAP on coronavirus epitopes from the IEDB (Methods).

To evaluate the accuracy of the trained model for our specific research question, we reserved all 66 experimentally validated 9- and 10-mer Wuhan-mutated pMHC and we compared their predicted scores with randomly sampled ‘negative’ sets ten times (66 per repetition, Methods). We demonstrated that TRAP could identify immunogenic Wuhan-mutated pMHC with a ROC-AUC of ∼0.76 (Fig 3B) and a precision-recall area-under-the-curve of ∼0.82 (S3A). Therefore, we determined the model was fit for purpose for predicting the impact of Omicron mutations on T cell recognition potential of Wuhan-mutated pMHC.

**Figure 3:**
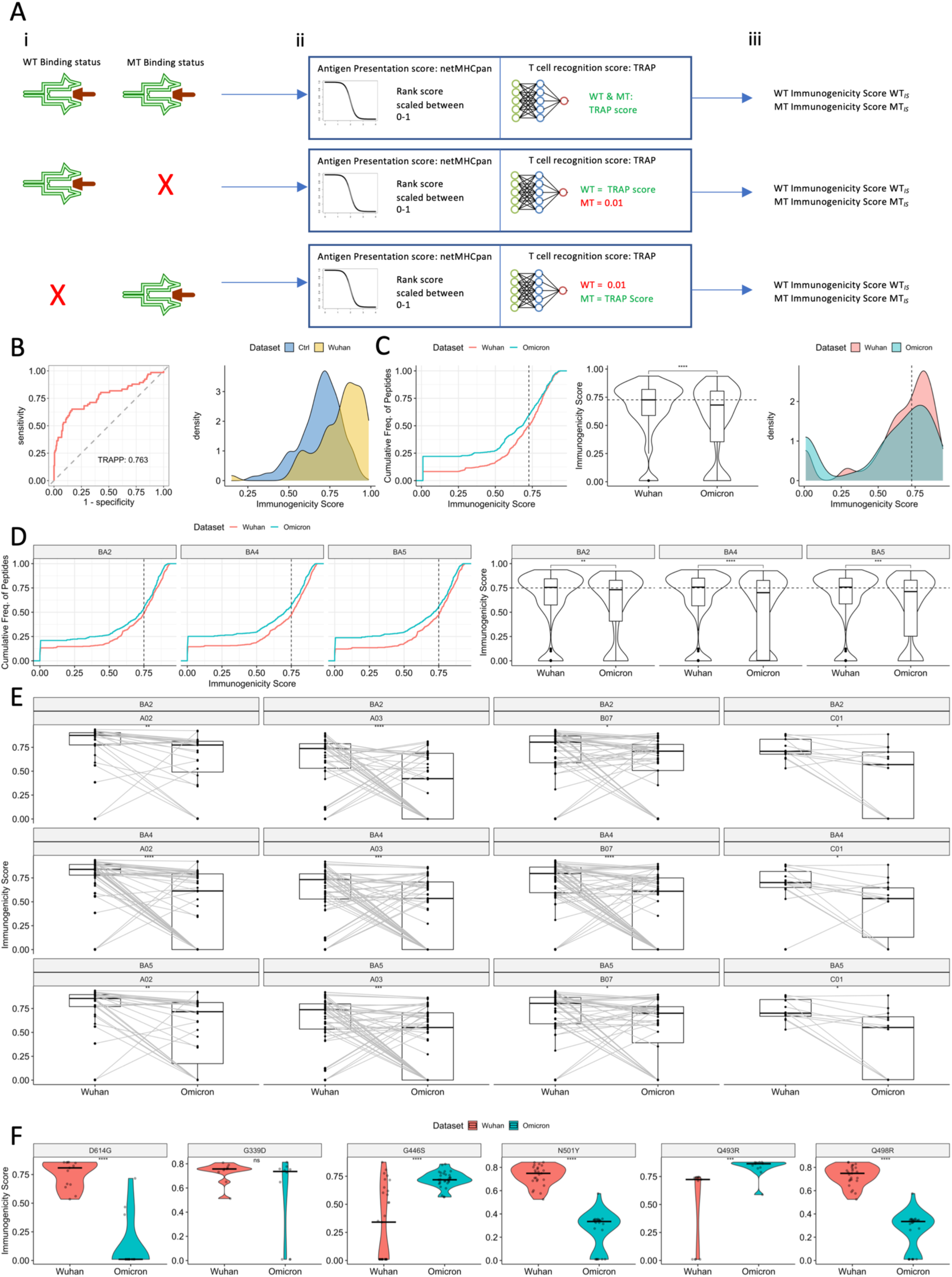
The predicted impact of BA1 Omicron and subvariant mutations on T cell immunogenicity of SARS-CoV-2 CD8+ T cell targets. Analyses were only performed for the set of Wuhan Hu-1 CD8+ T cell targets with a mutation and their mutated counterparts (WT-MT). T cell immunogenicity incorporates i) antigen presentation predictions by netMHCpan 4.1, and ii) T cell recognition predictions by TRAP. A) an overview of how the T cell immunogenicity scores were generated and the different scenarios by which imputations of pseudo-zero ‘T cell recognition’ scores (0.01) were made. TRAP by definition makes T cell recognition predictions against peptides bound to MHC *k*. Thus, given three potential bi-directional changes in WT vs. MT binding status, three scenarios are captured. Row 1 shows a setting where for a particular CD8+ T cell target, the wildtype peptide and mutant peptide, both bind MHC *k*. Here, antigen presentation predictions were made by netMHCpan which were combined with T cell recognition scores from TRAP (Methods for full details), no imputations were made. The immunogenicity score combines these two predictions. Row 2 shows a setting where the wildtype peptide binds MHC *k*, although the mutant – following Omicron mutation – does not. Here, antigen presentation is predicted using netMHCpan for both wild-type and mutant (the mutant will accordingly receive a low MHC binding score). TRAP however cannot make a T cell recognition prediction for a peptide not predicted to bind MHC *k*, therefore we impute a pseudo-zero value of 0.01 for the mutant. A similar situation is depicted in row 3, however, here, the wildtype was not predicted to bind MHC *k* thus was not immunogenic, however after Omicron mutation, antigen presentation status was observed. Here, we impute a TRAP score for the nonbinding wild type and make a TRAP prediction for the presented mutant. B) ROC-curve and density plot evaluating the performance of TRAP against 66 known SARS-CoV-2 Wuhan Hu-1 CD8+ T cell targets with a mutation in Omicron (immunogenic Wuhan set, yellow) vs. 10 sets of 66 functionally evaluated non-immunogenic SARS-CoV-2 prediction scores made by TRAP, which were randomly sampled from a 10-fold cross-validation of training data (control, blue). ROC-curves show the performance of a model through perturbing classification thresholding and visualizing the true positive rate (fraction of true positives/all true positives) against the false positive rate (fraction of false positives/all true negatives). Curve information is summarized using the area-under-the-curve. Given a balanced dataset for binary classification (50% per classification), an unskilled model will have a ROC-AUC of 0.5, reflecting only the balance in the dataset. A perfect model would have a ROC-AUC of 1.0. C) empirical cumulative distribution (ecdf) plot, violin plot and density plot comparing the predicted T cell immunogenicity scores of Wuhan-mutated vs. Omicron BA1 pMHC. Significance was assessed using a Wilcoxon rank test. D) ECDF and violin plots comparing T cell immunogenicity scores of Wuhan-mutated pMHC vs. Omicron subvariant BA2, BA4, BA5 counterparts. Significance was assessed using Wilcoxon rank tests. E) Paired boxplots for different HLA supertypes (HLA-A02, HLA-A03, HLA-B07, HLA-C01), comparing the T cell immunogenicity scores between Wuhan-mutated epitopes and Omicron variant mutated counterparts. Significance was assessed using paired Wilcoxon rank tests. F) Violin plots contrasting the Wuhan vs. BA1 Omicron T cell immunogenicity scores for spike mutations; D614G, G446S, G339D, N501Y, Q493R, Q498R. Significance was assessed using a Wilcoxon rank test.

To evaluate the extent that mutation affects T cell immunogenicity, we generated ‘immunogenicity’ scores by combining 1) a ‘netMHCpan’ antigen presentation score and 2) the ‘TRAP’ T cell recognition score (see Fig3A and Methods for complete details). By examining immunogenicity scores of 580 paired Wuhan-mutated WT vs. MT pMHC (from any protein), we observed that while global predicted T cell immunogenicity is preserved upon Omicron, BA1 epitopes show a subtle reduction when compared with their Wuhan Hu-1 counterparts (Fig 3C, Supplementary Datafile 1).

We repeated this analysis for Omicron subvariants BA2, BA4 and BA5 and found the same trend (Fig 3D, Supplementary Datafile 2). We also examined cross-HLA variation and observed reduced BA1 T cell immunogenicity for HLA-A02 and -C01 ligands (S3B). For BA2, BA4 and BA5 on the other hand we observed significant impairments in T cell immunogenicity for HLA-A02, -A03, -B07 and -C01 ligands (Fig 3E). Interestingly, the magnitude of impairment per supertype varied between subvariants, suggesting that different mutations across Omicron-based VOC produce nuanced effects on T cell immunogenicity that appear to be HLA-dependent.

We therefore next investigated how specific point mutations affect the immunogenicity of Wuhan vs. Omicron pMHC (and its subvariants). Interestingly, we observed substantial variation in the effects of mutation on BA1 T cell immunogenicity (Fig 3F, S3C). Indeed, mutations such as D614G, N501Y, Q498R are predicted to significantly reduce T cell immunogenicity, whereas mutations such as G446S[37] and Q493R may potentiate it (Fig 3F). In partial agreement with observations by Li Y et al[38], our modelling predicts that G339D impairs several pMHC, although we did not observe a significant reduction in T cell immunogenicity, suggesting variation regarding how this mutation affects different pMHC. Notably, Q→R substitutions in different positions of spike protein (P493 vs. P498) are predicted to lead to opposite effects, indicating that the same amino acid change at different positions produces opposite effects on T cell immunogenicity. By examining characteristic mutations of BA2, BA4 or BA5, we again observed distinct effects (S3D).

Lastly, we integrated our data with metadata of Naranbhai et al’s patients who exhibited impaired T cell responses. We found that for 6/10 patients, >=40% of their carried HLAs were associated with reductions in predicted T cell immunogenicity given BA1 mutated epitopes, while for two patients, >=60% of HLAs are impacted (S3E). Although we cannot draw a firm conclusion due to absence of control data, these findings build on the insight of Naranbhai et al., who argued that HLA genotype of these patients may in-part contribute to the impairments observed in their study.

Overall, our data indicates that while broadly the T cell response is preserved following Omicron mutations, their impact on the landscape of T cell immunogenicity is divergent and heterogeneous. This work provides insights into how different mutations are likely to affect T cell immunogenicity.

### Relative impact of Omicron mutation on potential and breadth of T cell responses

We have observed that Omicron mutations have disparate effects on pMHC immunogenicity. In general, mutation(s) at a CD8+ T cell target may impact the overall potential (i.e., change in likelihood of T cell response) and breadth (change in the number of cognate T cells in a given repertoire). To explore this further, we set out to i) profile the relative impact of Omicron mutation on likelihood of T cell immunogenicity (potential) for each pMHC (with respect to Wuhan Hu-1) and ii) estimate the effect of mutation on promiscuity of binding cognate TCRs (breadth). To achieve the first goal, we computed a metric which we termed ‘<u>r</u>elative immunogenic potential’ (RI) (Methods). RI is computed for each *pMHC* and a score of >0 indicates that the Omicron mutation(s) potentiates immunogenicity, while <0 indicates impairment (Supplementary Datafile 3).

At the same time, we aimed to estimate the effects of BA1 Omicron mutations on response breadth. A leading hypothesis for increased global rates of children with severe acute hepatitis, is that SARS-CoV-2 viral persistence may promote repeated immune activation. A motif within SARS-CoV-2 spike protein with resemblance to the SEB superantigen has been hypothesised to trigger broad, promiscuous T cell activation. As Omicron and its subvariants spread globally, we predicted whether Omicron mutations may alter the breadth of TCRs recognising mutated peptides.

We thus retrained TITAN, a state-of-the-art CNN model for computational mapping of TCRs to their target antigens (see Methods, Supplementary Methods). Therefore, TITAN allows interrogation of the breadth of T cell responses to a given epitope. After observing reasonable performance (ROC-AUC 0.76 albeit with considerable variation across epitopes), we took our unseen TCR dataset (162,930 TCRs, Supplementary Methods) and predicted their binding against each (unseen) Wuhan-mutated or counterpart Omicron peptides. TITAN does not consider HLA, thus, for each *peptide*, we calculated RTP: ‘<u>r</u>elative <u>T</u>CR <u>p</u>romiscuity (Methods),’ comparing the number of TCRs predicted to bind the Wuhan WT vs Omicron MT. RTP>0 reflects increased breadth after Omicron mutation; simply, binding more TCRs.

By analysing both ‘RI’ and ‘RTP’, we found a divergent landscape for the impact of BA1 mutations on immunogenic potential (i.e., RI, Fig. 4A) but no pattern for breadth (S4A). There are small to moderate impairments (RI <0) in potential for a substantial number of pMHC, while others are eliminated as CD8+ T cell targets through inability to bind MHC. In contrast, there are pMHC with enhanced potential following the mutation in Omicron. This analysis additionally suggests that whereas HLA presentation plays a permissive role in the likelihood of T cell immunogenicity, it does not seem to be contributing to the breadth.

**Figure 4:**
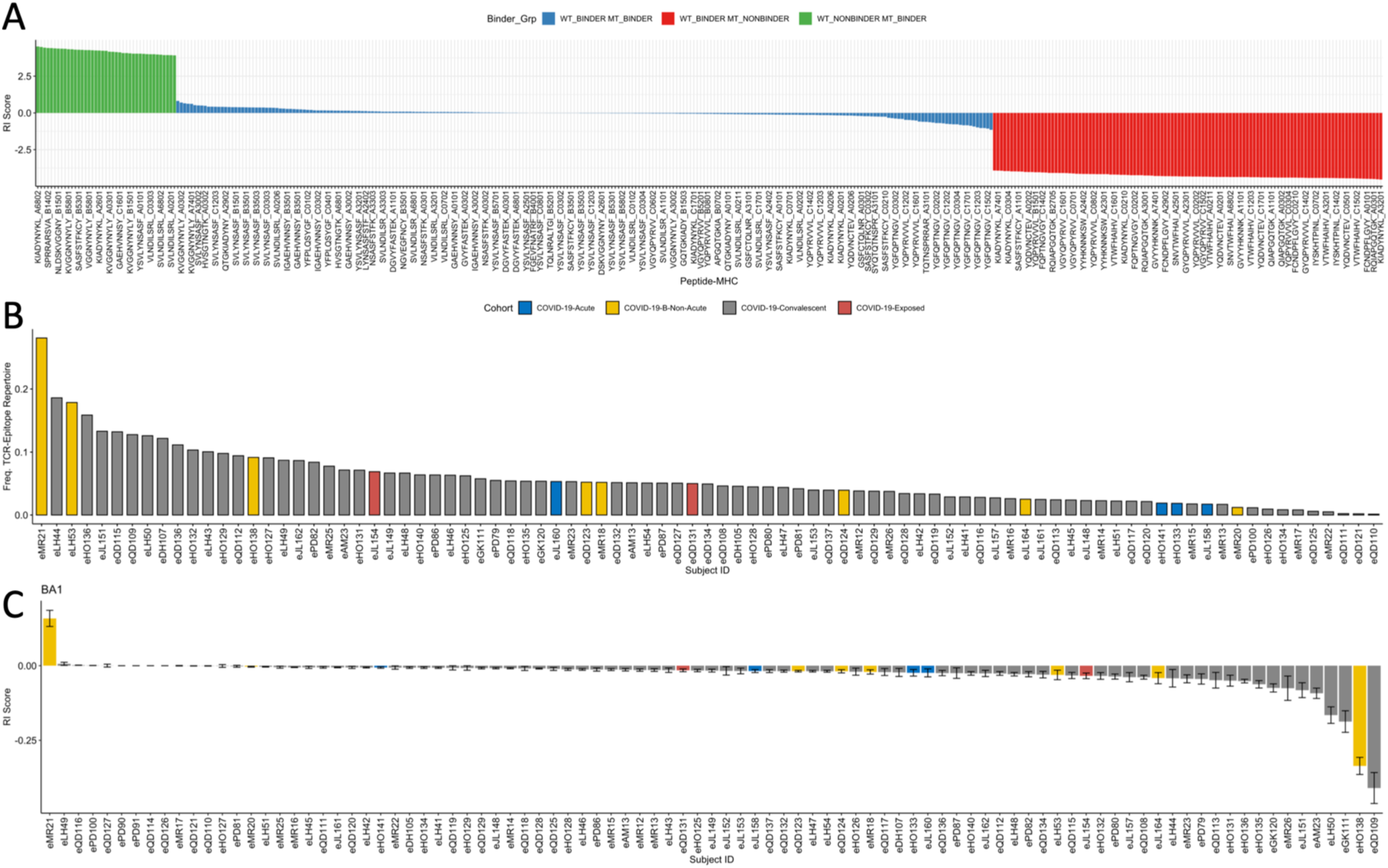
The overall impact of Omicron mutations on immunogenic potential of pMHC and individual TCR repertoires. A) Barplots showing the relative immunogenic potential (RI) score for each pMHC affected by a mutation in BA1 Omicron. Labels are truncated for visual clarity. RI scores for each pMHC are supplied in Supplementary Datafile 3. Color labelling represents groups of pMHC by wildtype←→ mutant binding status: WT_BINDER_MT_BINDER (blue), WT_BINDER_MT_NONBINDER (red), WT_NONBINDER_MT_BINDER (green), representing respectively pMHC where both the WT and MT bind the same MHC (blue), the WT binds a particular MHC but the mutant does not (red), the WT does not bind a particular MHC that the mutant does (green). RI is produced for each pMHC. B) Barplot showing the frequency of individual MIRA TCR-Epitope repertoires (excluding healthy individuals) which exhibit a mutation in BA1 Omicron. C) Barplot showing the mean +/− standard error for ‘RI’ scores for each individual MIRA TCR-Epitope repertoire. BA1 only. As MIRA antigen-specific data does not explicitly label the bound MHC to the antigen, RI scores here are computed for each peptide, thus represent the mean RI across predicted MHC (pan-HLA RI). Scores of ‘zero’ are imputed for non- summarised here for each patient.

While a handful of peptides that compose pMHC with enhanced potential are predicted to additionally bind more TCRs (Fig S4A/Fig. 4A), we were unable to draw meaningful conclusions regarding whether mutations impacted TCR breadth. Nevertheless, we propose that these pMHC with >0 RI score (enhanced potential) which are composed of peptides with predicted RTP >0 (increased breadth), may be of interest for investigations into post-COVID- 19 inflammatory disorders.

Thus far, our predictions show that mutations in Omicron BA1 (and its subvariants: S4B) produce heterogeneous effects on the immunogenic potential of pMHC, including some complexes that are removed as CD8+ T cell targets. Furthermore, we have observed that peptides bound to certain HLAs (e.g., A03, A02, B07) are associated with a reduction in immunogenicity, while other HLA are not. Collectively, this heterogeneous impact implies that individuals with certain HLAs and/or TCR repertoires biased toward certain epitopes will be more affected than others.

We thus sought to expand this approach to investigate the impact of mutations on antigen-specific T cell immunogenicity potential, at the level of individual TCR repertoires. We therefore extracted 160k high confidence SARS-CoV-2-specific TCRs from 93 COVID-19 patients (79 convalescent + 14 acute/non-acute/exposed) from the ‘MIRA’ dataset[39]. The data consist of 543 Wuhan Hu-1 SARS-CoV-2 peptides, 46 (∼8%) of which are mutated in Omicron. These individuals’ TCR repertoires were subjected to our RI score (see Methods).

We therefore treated each resulting TCR-epitope repertoire as a representation of Wuhan Hu- 1 induced memory and interrogated how Omicron mutations may affect overall potential of T cell response. First, for each subject we calculated the proportion of their TCR-epitope repertoire which targets Wuhan Hu-1 epitopes which exhibit a mutation in BA1 Omicron (Fig 4B). Consistent with our previous observations, we observed a variety of different impact, ranging from <1% to >20%. We next analysed the overall effects given the average RI scores for each individual’s antigen specific TCR repertoire (see Methods).

We found that while many individuals were likely to exhibit no effect given overall RI scores, a set of individuals showed impairments (Fig 4C, S4C). We did not find any association between these impaired individuals and available clinical metadata or e.g., size of the responding repertoires (S4D). It is plausible that reduced immunogenic potential that impacts overall T cell response, may in-part contribute to breakthrough infection, although further investigation is clearly required. Nevertheless, our data supports the hypothesis that an underlying cause of the heterogeneity observed with T cell impairment given VOC mutations is due to biased responses toward certain epitopes exhibiting impairments after mutation, which may be conditioned by HLA genotype.

### Inference of SARS-CoV-2 T cell escape potential by *in silico* mutagenesis

The set of VOC mutations that affect CD8+ T cell epitopes are a subset of theoretical mutational space; however, the underlying sampling mechanism is unclear. Sampling under some form of evolutionary pressure may suggest that the virus is taking a path to escape cellular immunity. Indeed, as optimal ACE2 affinity and cell entry are approached, wider forms of escape are likely to be evaluated by the virus[32]. A first step toward addressing this fundamental question and estimating the impact of emergent mutations on T cell responses, is to evaluate the impact of each theoretical single point mutation in a CD8+ T cell epitope on its immunogenicity.

To address this, we combined *in silico* mutagenesis and our immunogenicity modelling. As a proof of concept, we used the BA1 Wuhan-mutated 9/10-mer epitopes as a model system. For each Wuhan-mutated wild-type peptide (n=66) (WT_i_), we generated each theoretical mutant (MT_k=1‥n_, n=171 for 9-mers and 190 for 10-mers, see Methods). We then produced immunogenicity scores for predicted peptide-MHC as described previously (Supplementary Datafile 3), which we subsequently analysed in a pan-HLA manner (see Fig 3A and Methods).

Next, we analysed log ratios between each MT_k_ and its corresponding WT_i_ at TCR contact positions of 9-mers. Focusing on each amino acid (and not position), we found that the <u>removal</u> of polar residues cysteine (C), glycine (G), and basic lysine (K) from wildtype peptides increased immunogenicity (log ratio >0) (Fig 5A-left, S5A). Similarly, by incorporating these residues in the mutant (i.e., <u>replacement</u>), immunogenicity was reduced (Fig 5A-right, S5A). On the other hand, in most cases (6/8) the <u>removal</u> of hydrophobic residues (tryptophan (W), phenylalanine (F), alanine (A) etc) in TCR contact positions impaired immunogenicity. Next, we examined the impact of full mutations (A_x_B: A=removal, x=sequence position, B=replacement) occurring in the TCR contact positions of 9-mers. Focusing on the mutations with highest and lowest 5% of logs ratios, we again observed that primarily, the removal of hydrophobic residues impaired immunogenicity (Fig 5B). Interestingly, mutations that removed a basic amino acid predominately potentiated immunogenicity, while removals of phenylalanine (F) or tyrosine (Y) from P3 were commonly found amongst the most detrimental. Similarly, by examining TCR contact positions of 10-mers, removals of arginine (R), glycine (G) and lysine (K) potentiated immunogenicity, whereas removals of hydrophobic residues were again detrimental (Fig. 5C, S5B).

**Figure 5:**
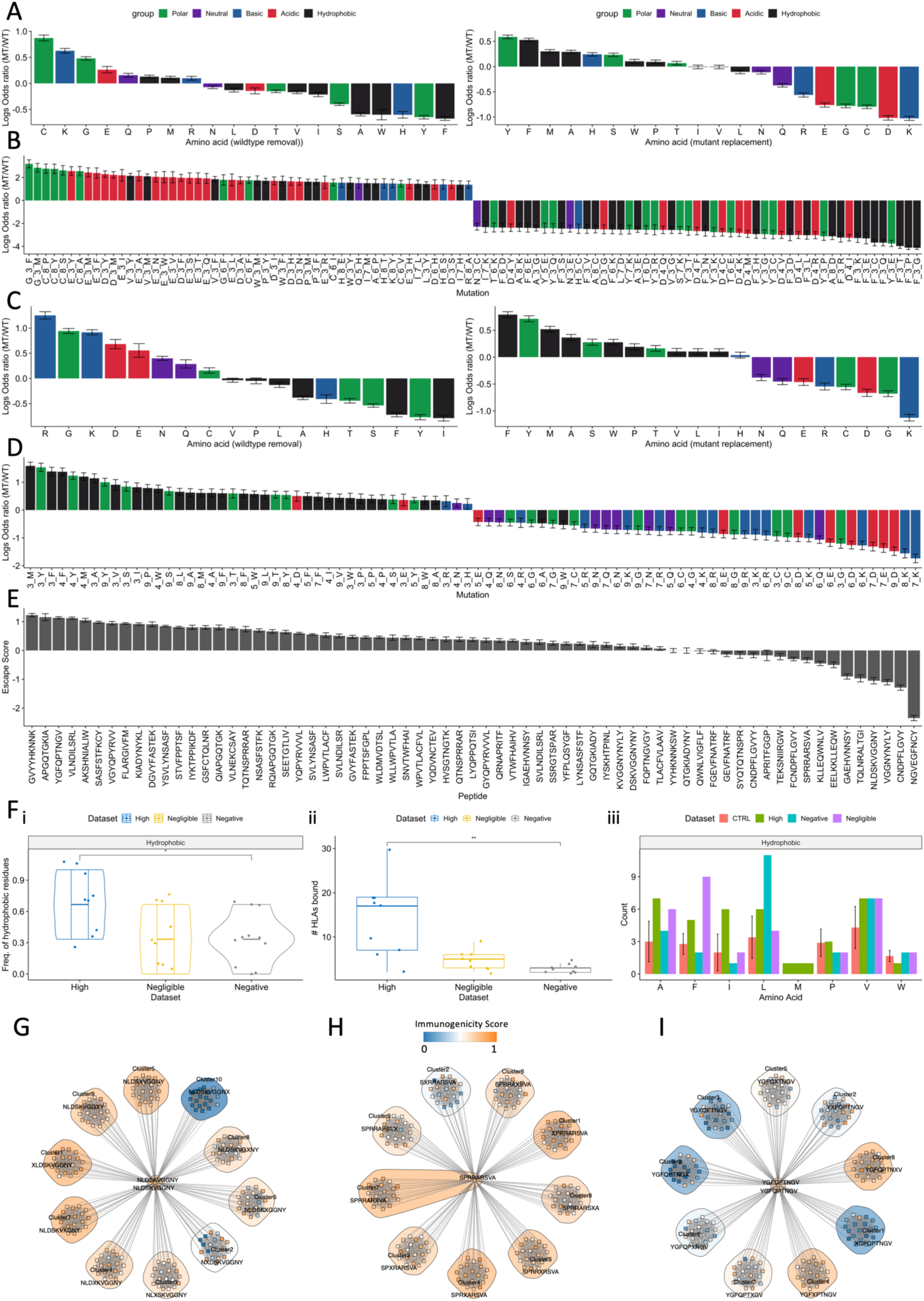
The effects of theoretical mutations on SARS-CoV-2-specific CD8+ T cell epitopes. A-D) Barplots showing the mean+/− standard error of log ratios derived from comparing mutant/wildtype immunogenicity scores from simulated mutations in TCR contact positions of A-B) 9-mers and C-D) 10-mers. Negative log ratios demonstrate that the mutant is less immunogenic than the wildtype and vice versa. A) Barplot showing for each amino acid residue, the mean+/− standard error across of all log ratios (9-mers). Left plot shows the log ratios upon removing a particular amino acid from the wildtype sequence. Right plot shows the log ratios upon imputing a particular amino acid in the mutant. Color labels show chemistry of amino acids. B) The highest and lowest 5% of log ratios derived by analysing the full mutations (A_x_B) simulated in TCR contact positions of 9mers. Color label shows the chemistry of the removed amino acid. C) Barplot showing for each amino acid residue, the mean+/− standard error across all log ratios (10- mers). Left plot shows the log ratios derived from removing a particular amino acid from the wildtype sequence. Right plot shows the log ratios derived from imputing a particular amino acid into the mutant. Color labels show chemistry of amino acids. D) Highest and lowest 30% of log ratios showing impact on immunogenicity for replacing residues (i.e., those amino acids inserted) into position X of TCR contact positions of 10-mers. E) Barplot showing ‘escape scores’: the mean +/− standard error of the difference between WT and MT (WT minus MT) immunogenicity scores affecting each assessed SARS-CoV-2 CD8+ T cell target (Methods for full details). F) i) Violin plot comparing the proportion of hydrophobic residues in anchor positions of three groups of peptides (n=9 per group, containing both 9- and 10-mers): blue) ‘high’, yellow) ‘negligible’ and grey) ‘negative’, grouping epitopes with blue) highest escape score, yellow) escape score approx. 0, and grey) negative escape scores, respectively. 9 samples were chosen to maximise samples whilst comparing the same quantity of peptides across groups. Significance was assessed using a Wilcoxon rank test. ii) Boxplots comparing the number of HLAs predicted to bind the 9 selected wildtype peptides for each ‘escape score’ group of interest. iii) Barplot showing the number of times each hydrophobic amino acid is observed in wildtype peptides composing the three ‘escape’ groups. A control group (CTRL, red) was generated to assess significance. The control group represents a bootstrapped background distribution of counts of hydrophobic amino acids from sampling 9 peptides randomly from the entire distribution of Wuhan-mutated 9- and 10-mer peptides. Crossbars show standard error. G-I) Neighbor network diagrams depicting the impact of mutation via network trajectories from the wildtype peptide (center) to each single amino acid variant. Clusters and individual mutants are color labelled by immunogenicity score. Groups are clustered by location of the substitution. An X in the consensus sequence by the cluster indicates the position of the substitution that characterises the cluster.

Interestingly for 10-mers in TCR contact positions, when we examined the impact of the *replacing* amino acid in position *x* we found a skew in underlying chemistry associated to immunogenicity enhancements or detriments (Fig 5D). In fact, hydrophobic and polar residues dominated the most immunogenic replacements, while detrimental replacements comprised more of a balance of acidic, basic and neutral residues. These findings not only provide a viewpoint into the molecular biology that underpins our immunogenicity modelling, but also suggest that physicochemical properties of amino acids (wildtype and/or replacement) may be critical for estimating the impact of mutation on T cell immunogenicity and contribute to how escape-prone a specific epitope is.

To identify the most and least escape-prone CD8+ T cell targets given theoretical mutations, we defined ‘escape score’ (Methods), as average log ratio changes across all mutants for each epitope of interest (Fig 5E, Supplementary Datafile 5). We found varying degrees of overall impact associated with three groups: 1) those predicted with *high* escape scores e.g., GVY* and YGF* showing overall impairment to immunogenicity, 2) those predicted with *negligible* escape scores, thus are tolerant to mutation (mean+/−se approx. 0) e.g., TLAC*, YYH*, and 3) those with a *negative* ‘escape score’ e.g., NGE*, NLD*, which are on the other hand predicted with global <u>improvements</u> to their immunogenicity as a result of mutation.

Given the associations between removal of hydrophobic residues and impaired immunogenicity, we explored whether properties of wildtype peptides may contribute to ‘escape score’. We found that escape-prone wildtype peptides have a slightly higher proportion of hydrophobic residues compared to those where mutation potentiated immunogenicity (Fig. 5F-i). Additionally, we found that escape-prone epitopes bind more HLA (Fig. 5F-ii), which may increase opportunities for escape. Furthermore, wildtype epitopes containing phenylalanine (F) seem to be more tolerant to mutation (Fig 5F-iii). These data suggest that while escape potential is multifaceted, the hydrophobic complexion and HLA promiscuity of the wildtype may guide the likelihood and direction that mutation impacts its immunogenicity.

Next, to visualise in detail the predicted mutational trajectories of each epitope, we implemented the Neighbour Network strategy from Ogishi et al[40] (Methods).

Through theoretical mutations, NLD* was predicted with overall improvements to its immunogenicity, thus a *negative* escape score (see Fig. 5E right). The wildtype of this epitope is already highly immunogenic; therefore, our analysis indicates that this epitope is a robust target, while on average mutations enhanced immunogenicity (Fig 5G). Notably, this epitope could tolerate many mutations in the P2 anchor position, whereas mutations in the P10 anchor position eliminated immunogenicity. Given this epitope’s general tolerance to mutation, it may be of interest e.g., for T cell therapies.

SPR* should be robustly immunogenic given our predictions (Fig 5H). Only mutations in P2 considerably impaired immunogenicity. With Omicron, this epitope is affected by a P→H mutation in P2, which drastically compromised binding to HLA-B07 alleles. These data indicate that in the ‘real-world’ - through VOC - SPR* has taken one of the few available escape routes. This finding is particularly interesting because SPR* is one of three epitopes that contains the hypothesised superantigen motif ‘PRRA’ which has been implicated in post-COVID-19 inflammatory disorders. Our analysis showed that after VOC mutation, SPR* will possess a narrower MHC binding repertoire. It is therefore plausible that an escape route has been taken – perhaps due to a highly immunogenic Wuhan Hu-1 wildtype - leaving only a single, highly potent pMHC (HLA-B*14:02-SPR*).

Our analysis predicts a high escape score for YGF* (Fig. 5E, left) via substantial numbers of escape routes (Fig. 5I). These data suggest that most mutations in anchor positions P1 (cluster 1) and P9 (cluster 9), as well as TCR contacting P3 (cluster 3) will damage immunogenicity. YGF* is affected by three Omicron mutations: G496S, Q498R and N501Y affecting P2, P4, P7 of the epitope respectively. Our immunogenicity modelling predicts that across each bound MHC, Omicron causes low immunogenicity for YGF* (immunogenicity scores 0.26-0.34, Supplementary Datafile 1). Indeed, the predicted effects of Omicron’s impact given three aggregate mutations may be unsurprising, as neighbour-network analysis indicates that YGF* was highly susceptible even to single-point mutations.

Overall, this analysis highlights the utility of applying *in silico* mutagenesis and immunogenicity modelling to examine the effect of theoretical mutations on T cell responses. We have observed varied effects of single point mutations on the immunogenicity of different epitopes. The ability to predict the impact of mutations on T cell immunogenicity would provide a powerful way to quickly understand the impact of an emergent VOC on a critical arm of the immune response. In a small step in this direction, we have developed an R script which, for a given mutation, amino acid removal or replacement, provides an estimate of the effect on T cell immunogenicity, given our *in-silico* mutagenesis data. This work therefore provides a gateway toward a strategy that given further validation and data, has the potential to reduce uncertainty associated with emergent VOCs and their impact on T cell responses.

## Discussion

The threat of a future VOC characterised by T cell evasion is an ongoing concern. As SARS-CoV-2 approaches optimal cell entry, other selection advantages, e.g., T cell escape, may be utilised to increase pathogenic fitness[32]. While extensive research has been undertaken to estimate effects of SARS-CoV-2 mutations on antibody recognition, mutational impact on T cell epitopes is poorly understood. Indeed, the ability to rapidly estimate the extent to which mutations impact T cell immunogenicity would reduce uncertainty upon emergence of novel VOC.

In this study, we have used *in silico* approaches i) to examine how mutations among Omicron VOCs affect SARS-CoV-2-specific CD8+ T cell responses and ii) to provide a preliminary framework for how theoretical mutations can impact SARS-CoV-2 epitopes. Our study is a step toward estimating the impact of mutation(s) on T cell immunogenicity upon the emergence of novel VOC. We demonstrate how T cell immunogenicity modelling can be used to not only dissect the impact of *existing* mutations on T cell responses, but also to forecast *theoretical* mutations that may be most detrimental for T cell immunity when combined with *in silico* mutagenesis. We envision that this work can support ongoing surveillance and forecasting efforts[41–43] and serve as a foundation of a knowledge base for determining how VOC may impact T cell responses upon their emergence. As model performance improves and more data are generated, the uncertainty regarding how emergent VOCs may affect T cell immunity could be reduced.

As Dolton et al. recently highlighted, ‘broad brush’ approaches have failed to uncover evidence of T cell escape following key variant mutations[10,32]. Indeed, during the early stages of Omicron’s rise to global dominance, the overwhelming consensus argued that Omicron does not significantly impair T cell responses [7,10,44–49]. Due to the diversity of HLA and breadth of epitopes, it had rightly been reasoned that widespread and complete T cell escape is unlikely at this stage of the pandemic [11,16,50,51].

In that light, *in silico* studies e.g., the work by Nersisyan et al[29] compared all theoretical HLA ligands across specific VOC and concluded that T cell responses to Omicron were likely to be maintained effectively. However, not all HLA ligands can invoke T cell responses [35,36]. Furthermore, studies such as those of Naranbhai et al[17] and Reynolds et al[18] observed considerable numbers of patients with impaired T cell responses to Omicron infection [52]. Indeed, while widespread and complete escape is unlikely, it is plausible that detrimental mutations could impair certain individuals e.g., with certain HLAs and narrow TCR repertoires targeting impacted pMHC. Thus, we hypothesised that some Omicron mutations may be more detrimental than the consensus, affecting the responses of some individuals more than others. This hypothesis was supported by *in silico* work by Pretti et al[31] indicating heterogeneous effects on binding HLA, given mutations among VOCs prior to Omicron.

We thus investigated CD8+ T cell targets with a mutation in Omicron and used predictive modelling to infer the effects of each mutation on T cell immunogenicity. Our work supports a body of literature that indicates overall, there exists a subtle reduction in T cell immunogenicity with mutated epitopes in Omicron VOCs, compared to Wuhan Hu-1. By examining mutated epitopes in detail, our study extends this paradigm, revealing a divergent landscape for how different CD8+ T cell targets are affected by Omicron-based mutation(s). In support of our hypothesis, we found that this heterogeneous landscape may produce a net impairment on some individuals’ T cell response against Omicron VOC, likely conditioned by bias toward certain epitopes in individual repertoires and patient HLA genotype. We speculate that these combined factors may lead to T cell escape in some individuals and perhaps breakthrough infections, although – as many epitopes are not mutated in VOCs – the extent to which mutated epitopes contribute to clinical outcome remains to be elucidated.

Evidence of SARS-CoV-2-specific T cell escape has begun to emerge. For example, Stavenich et al. [53] identified two mutations in a non-Hodgkin’s lymphoma patient infected with SARS-CoV-2, which escaped CD8+ T cell responses. Recently, Dolton et al. focused on the dominant HLA-A02 epitope YLQ, which was found to evade >175 TCRs due to a P→L mutation. Before Omicron’s emergence, Hamelin et al. [33] found that proline removals in Delta and Alpha could damage HLA-B07 pMHC binding. Extending these insights, our modelling predicts that P→X Omicron mutations eliminate certain HLA-B07 pMHC. Our work, combined with Hamelin and Dolton’s indicates that P→X mutations – which are now observed across Omicron variants that have infected large proportions of populations – in CD8+ T cell epitopes should be a priority for surveillance regarding T cell escape in HLA-B07 individuals and for associations with breakthrough infections. Overall, our data combined with recent studies is starting to indicate a more nuanced outlook regarding the impact of VOC mutations on T cell responses, than the consensus [17,31–33,52,54].

There are ongoing efforts to characterise disorders that are considered ‘post-COVID’ complications, ranging from so-called long-COVID [21,22,55–58] to broad inflammatory disease [20,23–25]. A leading hypothesis underpinning observations of multisystem inflammatory disorder and severe acute hepatitis, is that a theoretical SARS-CoV-2 superantigen promotes aberrant T cell activation [20,23]. The ‘PRRA’ motif, part of the hypothesised core of a SARS-CoV-2 superantigen [26,27] overlays three CD8+ T cell epitopes that have mutations in BA1 Omicron. Our data indicates that after Omicron mutation, 9/15 of the pMHC containing this theoretical superantigen core are removed as HLA ligands. We propose that remaining presenters HLA-A*33:03, -A31:03, -B14:02 be investigated for associations with inflammatory disorders following COVID-19 infection. More generally, we have predicted that a set of pMHC are likely to exhibit increased immunogenicity and bind more TCRs following Omicron mutation. Overall, further investigation is warranted to assess these particular CD8+ T cell targets, HLAs and cognate TCRs, for associations with post-COVID inflammatory disorders.

With potential exceptions (SPR*, Fig 5H), our data do not suggest that reductions in T cell immunogenicity observed with Omicron were the result of immune pressure. Nevertheless, many evaluated epitopes were predicted to be escape-prone, albeit with variation, given theoretical single point substitution. This insight could be concerning, given the considerable number of unexplored mutations available for SARS-CoV-2 that our data estimate could lead to T cell escape. Notwithstanding, mutations are not uniformly generated; thus, future work should seek to understand the likelihood of each potentially problematic theoretical mutation, perhaps by simulating evolution e.g., using SANTA-SIM[59] or forecasting driver mutations[43].

Our approach may have a limitation in calculating RI scores, where we modelled antigen presentation in a binary manner and imputed pseudo-zero T cell recognition scores for nonbinders. Here, if a WT peptide binds MHC *k* while the mutant does not, the immunogenicity equation (product of antigen presentation and T cell recognition) for the mutant is skewed to values close to zero and vice versa. This causes a large effect on RI scores (Fig 4A). Modelling antigen presentation in this manner can be justified by its binary nature, thus we applied a negative logistic function to generate values approaching 1 for strong binders and 0 for non- binders. Furthermore, nonbinders cannot canonically be recognised by T cells; thus, a pseudo-zero score for T cell recognition is logical.

Another limitation is that we are unable to directly analyse the effects of mutations in T cell epitopes on breakthrough infections. This is due to the lack of publicly available data to support this analysis, which would require antigen-specific memory TCR repertoires linked with clinical information prior to and after Omicron infection. It will be important to examine whether patients who experienced breakthrough infections are enriched for T cell responses that are dominant toward epitopes we have predicted will exhibit a reduction or loss of immunogenicity after Omicron. This work would help us understand to what extent the loss of immunogenic epitopes due to mutation affects clinical outcomes.

In conclusion, we have utilised *in-silico* approaches to assess the impact of existing and theoretical mutations on SARS-CoV-2 CD8+ T cell targets. We reveal a divergent, heterogeneous landscape of impact for Omicron VOC. We proposed a framework for forecasting the effects of theoretical SARS-CoV-2 mutations using immunogenicity modelling and *in-silico* mutagenesis. We hope that our work provides a gateway toward a comprehensive and publicly available approach for assessing the impact of mutations on T cell immunogenicity upon emergence of novel VOC. In the case of detrimental impacts, we envision this approach, given further development, could rapidly generate a profile of affected epitopes and HLAs to estimate populations that are most likely be affected by novel VOC. Our framework should next be applied to all known SARS-CoV-2 CD8+ T cell epitopes to i) fully characterise escape-prone vs. robust epitopes, ii) identify further theoretical mutations warranting surveillance and iii) further molecular insight into the biology underpinning mutation impact.

## Methods

### Data Analysis

Data processing and analysis were performed with R or Python 3.7. Visualisations were made using R library *ggplot2* or *ggpubr*.

### Curating immunogenic SARS-CoV-2 CD8+ T cell targets

1406 unique immunogenic MHC-I SARS-CoV-2 epitope data evaluated in the context of humans were gathered from IEDB and MIRA. 26 of these epitopes were excluded due to origin in a non-ancestral strain. 1380 unique SARS-CoV-2 Wuhan Hu-1 immunogenic epitopes were selected for analysis.

### Retrieval of SARS-CoV-2 proteomes

The Wuhan Hu-1 proteome was obtained from: https://www.ncbi.nlm.nih.gov/nuccore/nc_045512.2 Representative strains for the proteomes of Omicron BA1, BA2, BA4 and BA5 were retrieved from BV-BRC: the SARS-CoV-2 Variants and Lineages of Concern resource: https://www.bv-brc.org/view/VariantLineage/#view_tab=overview.

### MHC Presentation Prediction

Antigen presentation by MHC class I was predicted using NetMHCpan v4.1 against 64 HLA types. These HLA are listed in Supplementary Methods. Peptide-MHCs with a binding affinity rank score <= 2.0 were classified as binders.

### Immunogenic Wuhan Hu-1 epitopes with a mutation in Omicron and its subvariants

For each Wuhan Hu-1 epitope, we performed a local-global alignment using *pairwiseAlignment* function from the R package *Biostrings* against each protein in Omicron proteomes. The maximum alignment across the proteome was selected as the counterpart variant. Mutation information was recorded and mapped with a known list of mutations of the respective variant, downloaded from https://covariants.org/ and https://github.com/hodcroftlab/covariants.

### Mutation effect on antigen presentation

Paired WT-MT samples were analysed by comparing predicted binding affinity nM, netMHCpan rank and the ratio of mutant/wild-type predicted binding affinity predicted in nM (agretopicity). We focused on samples where *either* or *both* the WT and MT were predicted to bind a particular allele. The reasoning for this, was to incorporate the following three scenarios: i) where the WT is predicted to bind but the MT is not, ii) where the WT is not predicted to bind MHC but after mutation, the mutant becomes a binder, iii) where both the WT and MT bind a particular MHC.

### Training TRAP to predict immunogenic SARS-CoV-2 epitopes

Immunogenic and non-immunogenic peptides of length 9 and 10 were retrieved from the IEDB in July 2022. Peptide-MHC samples from antigen organisms containing the words “coronavirus” or “SARS” were retained. Binary ‘positive’ or ‘negative’ labels were extracted. 37 peptides with 1 immunogenic but >2 non-immunogenic observations were excluded as this suggests the immunogenic observation could not be replicated. Next, we confirmed that all epitopes only contain amino acid characters. We excluded any pMHC that did not have four-digit resolution HLA information. These filters left 1641 distinct pMHC for analysis; 566 (34.5%) were non-immunogenic, 1075 (65.5%) were immunogenic.

For any remaining contradictory pMHC (those with both immunogenic and non-immunogenic observation(s)), we assumed these epitopes can be immunogenic thus we excluded the ‘negative’ observations. To avoid overfitting, we excluded any pMHC which contained Wuhan Hu-1 epitopes with a mutation in Omicron. We therefore used 1,511 coronavirus epitopes to train the coronavirus TRAP model, 1074 (∼71%) were immunogenic, 437 (∼29%) non-immunogenic. The TRAP model predicts CD8+ T cell recognition potential of HLA-I ligands from peptide sequences represented in ProtT5-XL-UniRef embeddings, MHC binding rank score predicted by netMHCpan and the proportion of hydrophobic amino acids using a 1D CNN architecture. Kernels are used in this model to extract recognition motifs from TCR contact positions.

### Evaluating TRAP’s performance

66 Functionally validated pMHC containing peptides with a mutation in Omicron were excluded from training and reserved for testing. To generate a ‘negative’ set, 66 predictions for non-immunogenic pMHC were sampled 10 times from a 10-fold cross-validation of the training data. At each sampling, ROC-AUC, PR-AUC, precision, recall, sensitivity, specificity, balanced accuracy, detection rate and prevalence, F1 and negative/positive predictive values were evaluated. For ROC-AUC and PR-AUC plots, the data across ten samplings were summarised.

### Comparing T cell immunogenicity between Wuhan Hu-1 and Omicron variants

We compared overall predicted T cell immunogenicity of Wuhan-mutated epitopes, and their mutant counterparts. We reasoned that ‘overall’ T cell immunogenicity encompasses two components: 1) antigen presentation and 2) T cell recognition, where antigen presentation is a pre-requisite for T cell recognition. We combined two metrics for 1) antigen presentation and 2) T cell recognition to generate an ‘overall’ immunogenicity score.

For antigen presentation, we generated netMHCpan percentile rank binding-affinity estimates for each peptide-MHC. As different HLA bind their ligands in different nM ranges, percentile rank was used to reduce bias across HLA alleles. Percentile rank was transformed between 0-1 by use of a negative logistic function:

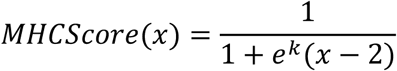

The function provides a value approaching 0 for a percentile rank much higher than 2.0, the recommended threshold for discriminating netMHCpan binding. Where percentile rank = x=2, the function evaluates to 0.5. For very low percentile ranks, i.e., strong binders, the function gives a value approaching 1. K is a parameter which defines the shape of the curve. We selected k=1.5, to avoid too strong skews either side of the midpoint percentile ranks. A range of values for *k* were explored with no change to conclusions. ‘MHCScore’ was generated for each WT and each MT pMHC of interest.

We generated ‘T cell recognition’ score for each WT and each MT pMHC using TRAP. As TRAP predicts T cell recognition, it naturally only considers samples in which binding to MHC is achieved. However, to address changes between WT and MT viral strains fairly, one must account for three groups of WT-MT MHC binding relationships: 1) samples where WT was predicted to bind MHC-*i*, but MT does not, 2) samples where WT and MT are both predicted to bind MHC-*i*, 3) samples where the WT was not predicted to bind MHC-*i* but MT is now predicted to bind. Samples where both WT and MT were not predicted to bind MHC were discarded.

In scenarios 1 and 3, TRAP cannot produce a score for both WT and MT pMHC, as either the WT or MT is not predicted to bind MHC-*i*. Therefore, for predicted nonbinders, we imputed a pseudo-zero ‘T cell recognition’ score of 0.01, as by definition nonbinders cannot be recognised by T cells. Any binders were subjected to TRAP to generate a ‘T cell recognition’ score.

The ‘overall’ immunogenicity score for each pMHC of interest, is given by:

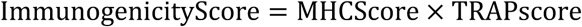

Samples in which both WT pMHC and MT pMHC are predicted with immunogenicity scores <0.35 were excluded, as we estimate that these are not immunogenic and therefore are not of interest.

### Relative Immunogenic potential (RI score)

For a pMHC of interest:

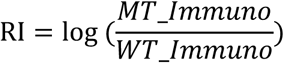

where *MT_Immuno, WT_Immuno* are the immunogenicity scores for the MT pMHC and the WT pMHC respectively. This score was generated for each pMHC complex assessed.

### Training TITAN to examine TCR-Epitope specificity

TITAN is a bimodal convolutional neural network[60] that encodes TCR and epitope sequences to predict the binding probability of TCR-epitope interactions. A limitation of TITAN is that it does not consider the HLA allele presenting an epitope. Therefore, predictions made using this model are interpreted and analysed in a ‘pan-peptide’ fashion.

For our work, we trained TITAN using the full TCR sequence: V, J gene sequences and CDR3. TCRs were encoded using BLOSUM62. Due to superior performance in the authors’ original study, we encoded the peptides using SMILES, which is an atom-level description commonly used in chemoinformatics. Full retraining details are provided in the Supplementary Methods.

### Relative TCR Promiscuity score (RTP)

TITAN was used to predict binding between epitope and TCR.

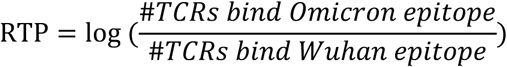

TITAN does not consider MHC. Therefore, RTP is computed for each peptide. RTP>0 reflects increased breadth after Omicron mutation; simply, binding more TCRs. RTP should be considered as a ‘pan peptide’ score, while RI is peptide-MHC specific.

### Analysis of the MIRA dataset

The COVID-19 MIRA dataset was downloaded from https://clients.adaptivebiotech.com/pub/covid-2020 with sample metadata. These data contain TCR repertoire data mapped to SARS-CoV-2 epitopes from patient cohorts. As our analysis focused on how Omicron may affect T cell memory, healthy patients were excluded. Given our focus on CD8+ T cells, class II TCR-epitope samples were excluded. Unproductive TCRs were excluded. We define ‘TCR-Epitope’ repertoire, as each subject’s collection of unique TCR-Epitope samples from their full TCR repertoire. While the MHC of each subject is often known, the MIRA dataset maps TCR-Epitope and does not explicitly link TCR-Epitope-MHC. In effect, by using these data, it is not always possible to infer the specific MHC linked to the specific TCR-Epitope sample. Therefore, we analysed the MIRA data in a ‘pan-peptide’ manner. To do this, for each peptide, we took the average RI score across predicted bound MHC after Omicron, generating a single score for each immunogenic peptide with a mutation in Omicron. These data were then integrated with the MIRA TCR repertoires. Any peptides in the repertoires that are not mutated in Omicron variant analysed were imputed with a ‘0’ RI score, reflecting no change.

### In-silico mutagenesis

As TRAP is limited to 9- and 10-mers, we focused on single-point substitutions only. Immunogenicity scores for each wild-type peptide *i* (original Wuhan Hu-1 epitope) and its theoretical mutants (MT_*k*_ ‥ MT_*n*_ where n=171 for 9-mers and 190 for 10-mers) against 64 HLA were generated as described previously.

For panels 5A-D, we analysed log ratios in a ‘pan-HLA’ manner. Thus, for each MT_k_, we then took the mean score across filtered MHC as their ‘immunogenicity score’. So, for each wild-type peptide *i*, we compared one score (its average across MHC), with *n* (n=171 for 9mers and 190 for 10mers) MT scores (each representing their average across MHC).

To generate the ‘escape score’, we took the mean +/−standard error of log ratios (MT/WT) across all substitutions and assessed MHC involving a particular epitope. The escape score was then multiplied by −1 to produce a score where >0 reflects escape opportunity. Therefore:

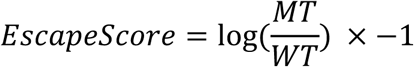

### Neighbor Networks

We adopted the approach and adapted the code from Ogishi et al. and their Repitope package: https://github.com/masato-ogishi/Repitope. We generated a network-style representation of the single amino acid substitution trajectories, where pairs of peptide sequences with just one alteration are defined as neighbors and regarded as edges. Clustering was performed using a walk-trap algorithm from the R *igraph* package. We adapted the Repitope code for our own use-case and to incorporate our own immunogenicity scores rather than those output from Repitope.

## Acknowledgements

HK was funded by UK MRC. This work was supported from the UK Medical Research Council with grant MC_UU_12010/3.

We thank D. Hudson for discussions regarding the manuscript, particularly surrounding TCR-epitope binding predictions.

## Author Contributions

HK supervised the project. HK and PRB conceived and designed the project. PRB performed computational analyses, with input and insights from HK, CHL, AA. CHL developed and ran the TRAP model. HK and PRB interpreted the results with insights from CL, AA. HK, AS funded the project. PRB, HK wrote the manuscript with contributions from CL, AA, AS.

## Data Availability Statement

Key datasets generated or analyzed during this study are included in this published article and its supplementary information files. Code used to perform the analysis can be found on github: [Enter on publication].

## References

1. Sigal A. Milder disease with Omicron: is it the virus or the pre-existing immunity? Nat Rev Immunol 2022; 22:69–71

2. Willett BJ, Grove J, Maclean OA, et al. The hyper-transmissible SARS-CoV-2 Omicron variant exhibits significant antigenic change, vaccine escape and a switch in cell entry mechanism. medRxiv 2022; 2022.01.03.21268111

3. Cele S, Jackson L, Khoury DS, et al. Omicron extensively but incompletely escapes Pfizer BNT162b2 neutralization. Nature 2022; 602:654–656

4. Cao Y, Wang J, Jian F, et al. Omicron escapes the majority of existing SARS-CoV-2 neutralizing antibodies. Nature 2022; 602:657–663

5. Cameroni E, Bowen JE, Rosen LE, et al. Broadly neutralizing antibodies overcome SARS-CoV-2 Omicron antigenic shift. Nature 2022; 602:664–670

6. Khan K, Karim F, Ganga Y, et al. Omicron BA.4/BA.5 escape neutralizing immunity elicited by BA.1 infection. Nat Commun 2022; 13:4686

7. Gao Y, Cai C, Grifoni A, et al. Ancestral SARS-CoV-2-specific T cells cross-recognize the Omicron variant. Nat Med 2022; 28:472–476

8. Grifoni A, Sette A. From Alpha to omicron: The response of T cells. Current Research in Immunology 2022; 3:146–150

9. Mclean G, Kamil J, Lee B, et al. The Impact of Evolving SARS-CoV-2 Mutations and Variants on COVID-19 Vaccines. 2022;

10. Tarke A, Sidney J, Methot N, et al. Impact of SARS-CoV-2 variants on the total CD4+ and CD8+ T cell reactivity in infected or vaccinated individuals. Cell Rep Med 2021; 2:

11. Alison Tarke A, Sidney J, Methot N, et al. Negligible impact of SARS-CoV-2 variants on CD4 + and CD8 + T cell reactivity in COVID-19 exposed donors and vaccinees.

12. Keeton R, Richardson SI, Moyo-Gwete T, et al. Prior infection with SARS-CoV-2 boosts and broadens Ad26.COV2.S immunogenicity in a variant-dependent manner. Cell Host Microbe 2021; 29:1611–1619.e5

13. Melo-González F, Soto JA, González LA, et al. Recognition of Variants of Concern by Antibodies and T Cells Induced by a SARS-CoV-2 Inactivated Vaccine. Front Immunol 2021; 12:4679

14. Riou C, Keeton R, Moyo-Gwete T, et al. Escape from recognition of SARS-CoV-2 variant spike epitopes but overall preservation of T cell immunity. Sci Transl Med 2022; 14:6824

15. Geers D, Shamier MC, Bogers S, et al. SARS-CoV-2 variants of concern partially escape humoral but not T-cell responses in COVID-19 convalescent donors and vaccinees. Sci Immunol 2021; 6:1750

16. Tarke A, Coelho CH, Zhang Z, et al. SARS-CoV-2 vaccination induces immunological T cell memory able to cross-recognize variants from Alpha to Omicron. Cell 2022; 185:847–859.e11

17. Naranbhai V, Nathan A, Kaseke C, et al. T cell reactivity to the SARS-CoV-2 Omicron variant is preserved in most but not all individuals. Cell 2022;

18. Reynolds CJ, Pade C, Gibbons JM, et al. Immune boosting by B.1.1.529 (Omicron) depends on previous SARS-CoV-2 exposure. Science 2022; eabq1841

19. Suryawanshi R, Ott M. SARS-CoV-2 hybrid immunity: silver bullet or silver lining? Nat Rev Immunol 2022; 1–2

20. Noval Rivas M, Porritt RA, Cheng MH, et al. Multisystem Inflammatory Syndrome in Children and Long COVID: The SARS-CoV-2 Viral Superantigen Hypothesis. Front Immunol 2022; 13:

21. Klein J, Wood J, Jaycox J, et al. Distinguishing features of Long COVID identified through immune profiling. medRxiv 2022; 2022.08.09.22278592

22. Al-Aly Z, Bowe B, Xie Y. Long COVID after breakthrough SARS-CoV-2 infection. Nat Med 2022; 1–7

23. Brodin P, Arditi M. Severe acute hepatitis in children: investigate SARS-CoV-2 superantigens. Lancet Gastroenterol Hepatol 2022; 0:

24. Sacco K, Castagnoli R, Vakkilainen S, et al. Immunopathological signatures in multisystem inflammatory syndrome in children and pediatric COVID-19. Nat Med 2022; 28:1050–1062

25. Porritt RA, Paschold L, Rivas MN, et al. HLA class I–associated expansion of TRBV11-2 T cells in multisystem inflammatory syndrome in children. Journal of Clinical Investigation 2021; 131:

26. Hamdy A, Leonardi A. Superantigens and SARS-CoV-2. Pathogens 2022; 11:390

27. Hongying Cheng M, Zhang S, Porritt RA, et al. Superantigenic character of an insert unique to SARS-CoV-2 spike supported by skewed TCR repertoire in patients with hyperinflammation.

28. Shen XR, Geng R, Li Q, et al. ACE2-independent infection of T lymphocytes by SARS-CoV-2. Signal Transduct Target Ther 2022; 7:1–11

29. Nersisyan S, Zhiyanov A, Zakharova M, et al. Alterations in SARS-CoV-2 Omicron and Delta peptides presentation by HLA molecules. PeerJ 2022; 10:e13354

30. Foix A, López D, Díez-Fuertes F, et al. Predicted impact of the viral mutational landscape on the cytotoxic response against SARS-CoV-2. PLoS Comput Biol 2022; 18:e1009726

31. Pretti MAM, Galvani RG, Scherer NM, et al. In silico analysis of mutant epitopes in new SARS-CoV-2 lineages suggest global enhanced CD8+ T cell reactivity and also signs of immune response escape. Infection, Genetics and Evolution 2022; 99:105236

32. Dolton G, Rius C, Hasan MS, et al. Emergence of immune escape at dominant SARS-CoV-2 killer T cell epitope. Cell 2022; 185:2936–2951.e19

33. Hamelin DJ, Fournelle D, Grenier JC, et al. The mutational landscape of SARS-CoV-2 variants diversifies T cell targets in an HLA-supertype-dependent manner. Cell Syst 2022; 13:143–157.e3

34. dos Santos Francisco R, Buhler S, Nunes JM, et al. HLA supertype variation across populations: new insights into the role of natural selection in the evolution of HLA-A and HLA-B polymorphisms. Immunogenetics 2015; 67:651–663

35. H Lee C, Antanaviciute A, R Buckley P, et al. To what extent does MHC binding translate to immunogenicity in humans? ImmunoInformatics 2021; 3–4:100006

36. Buckley PR, Lee CH, Ma R, et al. Evaluating performance of existing computational models in predicting CD8+ T cell pathogenic epitopes and cancer neoantigens. Brief Bioinform 2022; 23:1–18

37. Motozono C, Toyoda M, Tan TS, et al. The SARS-CoV-2 Omicron BA.1 spike G446S mutation potentiates antiviral T-cell recognition. Nat Commun 2022; 13:1–11

38. Li Y, Wang X, Jin J, et al. T-cell responses to SARS-CoV-2 Omicron spike epitopes with mutations after the third booster dose of an inactivated vaccine. J Med Virol 2022; 94:3998–4004

39. Nolan S, Vignali M, Klinger M, et al. A large-scale database of T-cell receptor beta (TCRβ) sequences and binding associations from natural and synthetic exposure to SARS-CoV-2. Res Sq 2020; 1–28

40. Ogishi M, Yotsuyanagi H. Quantitative prediction of the landscape of T cell epitope immunogenicity in sequence space. Front Immunol 2019; 10:827

41. DeGrace MM, Ghedin E, Frieman MB, et al. Defining the risk of SARS-CoV-2 variants on immune protection. Nature 2022; 605:640–652

42. Babady NE, Burckhardt RM, Krammer F, et al. Building a Resilient Scientific Network for COVID-19 and Beyond. 2022;

43. Maher MC, Bartha I, Weaver S, et al. Predicting the mutational drivers of future SARS-CoV-2 variants of concern. Sci Transl Med 2022; 14:3445

44. Flemming A. Omicron, the great escape artist. Nat Rev Immunol 2022; 22:75–75

45. Keeton R, Tincho MB, Ngomti A, et al. T cell responses to SARS-CoV-2 spike cross-recognize Omicron. 2022;

46. Choi SJ, Kim D-U, Noh JY, et al. T cell epitopes in SARS-CoV-2 proteins are substantially conserved in the Omicron variant. Cell Mol Immunol 2022; 1–2

47. Mazzoni A, Vanni A, Spinicci M, et al. SARS-CoV-2 Spike-Specific CD4+ T Cell Response Is Conserved Against Variants of Concern, Including Omicron. Front Immunol 2022; 13:121

48. Tarke A, Coelho CH, Zhang Z, et al. SARS-CoV-2 vaccination induces immunological T cell memory able to cross-recognize variants from Alpha to Omicron. Cell 2022; 0:

49. Martinez-Sobrido L, Almazan Toral F, Faraz Ahmed S, et al. SARS-CoV-2 T Cell Responses Elicited by COVID-19 Vaccines or Infection Are Expected to Remain Robust against Omicron. Viruses 2022, Vol. 14, Page 79 2022; 14:79

50. Grifoni A, Sidney J, Vita R, et al. SARS-CoV-2 Human T cell Epitopes: adaptive immune response against COVID-19. Cell Host Microbe 2021;

51. Grifoni A, Weiskopf D, Ramirez SI, et al. Targets of T Cell Responses to SARS-CoV-2 Coronavirus in Humans with COVID-19 Disease and Unexposed Individuals. Cell 2020; 181:1489–1501.e15

52. Kedzierska K, Thomas PG. Count on us: T-cells in SARS-CoV-2 infection and vaccination. Cell Rep Med 2022; 100562

53. Stanevich O, Alekseeva E, Sergeeva M, et al. SARS-CoV-2 escape from cytotoxic T cells during long-term COVID-19. Res Sq 2022;

54. Yu F, Tai W, Cheng G. T-cell immunity: a barrier to Omicron immune evasion. Signal Transduct Target Ther 2022; 7:1–3

55. Alwan NA. Lessons from Long COVID: working with patients to design better research. Nat Rev Immunol 2022; 22:201–202

56. Galán M, Vigón L, Fuertes D, et al. Persistent Overactive Cytotoxic Immune Response in a Spanish Cohort of Individuals With Long-COVID: Identification of Diagnostic Biomarkers. Front Immunol 2022; 0:1129

57. Szabo PA, Dogra P, Gray JI, et al. Longitudinal profiling of respiratory and systemic immune responses reveals myeloid cell-driven lung inflammation in severe COVID-19. Immunity 2021; 54:797–814.e6

58. Liu X, Zhu A, He J, et al. Single-cell analysis reveals macrophage-driven T cell dysfunction in severe COVID-19 patients. medRxiv 2020;

59. Jariani A, Warth C, Deforche K, et al. SANTA-SIM: simulating viral sequence evolution dynamics under selection and recombination. Virus Evol 2019; 5:

60. Weber A, Born J, Rodriguez Martínez M. TITAN: T-cell receptor specificity prediction with bimodal attention networks. Bioinformatics 2021; 37:i237–i244

