## Supplemental Figures for "A systems approach evaluating the impact of SARS-CoV-2 variant of concern mutations on CD8+ T cell responses"

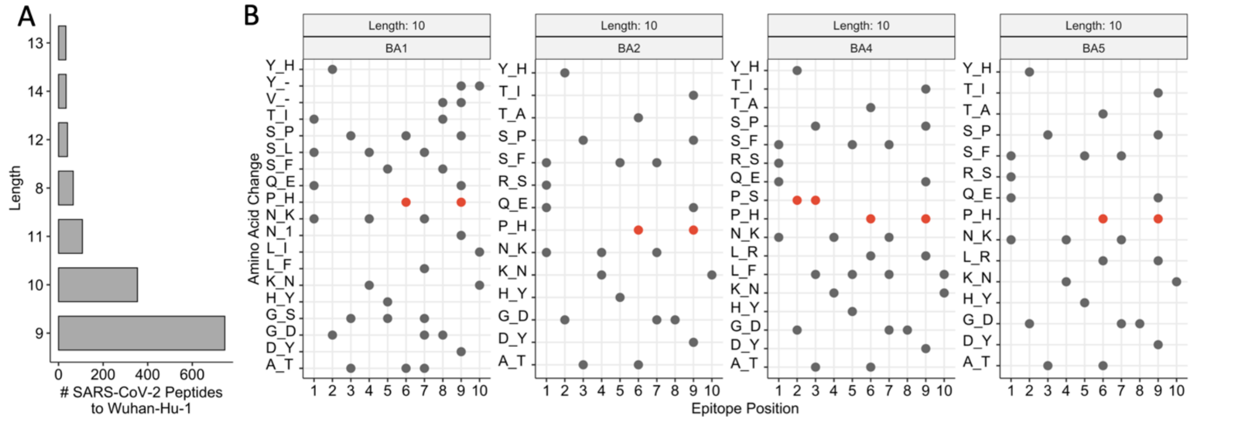


Supplementary Figure 1: A) The distribution of lengths of the unique SARS-CoV-2 ancestral Wuhan Hu-1 CD8+ T cell targets. B) The landscape of amino acid substitutions across different variants of concern within CD8+ T cell targets of length 10 residues. 'X_Y' on the y-axis indicates the removal of amino acid X, which is replaced by amino acid Y. The x-axis shows the position in the epitope that the mutation is observed. Red points highlight amino acid alterations which remove a 'Proline'.

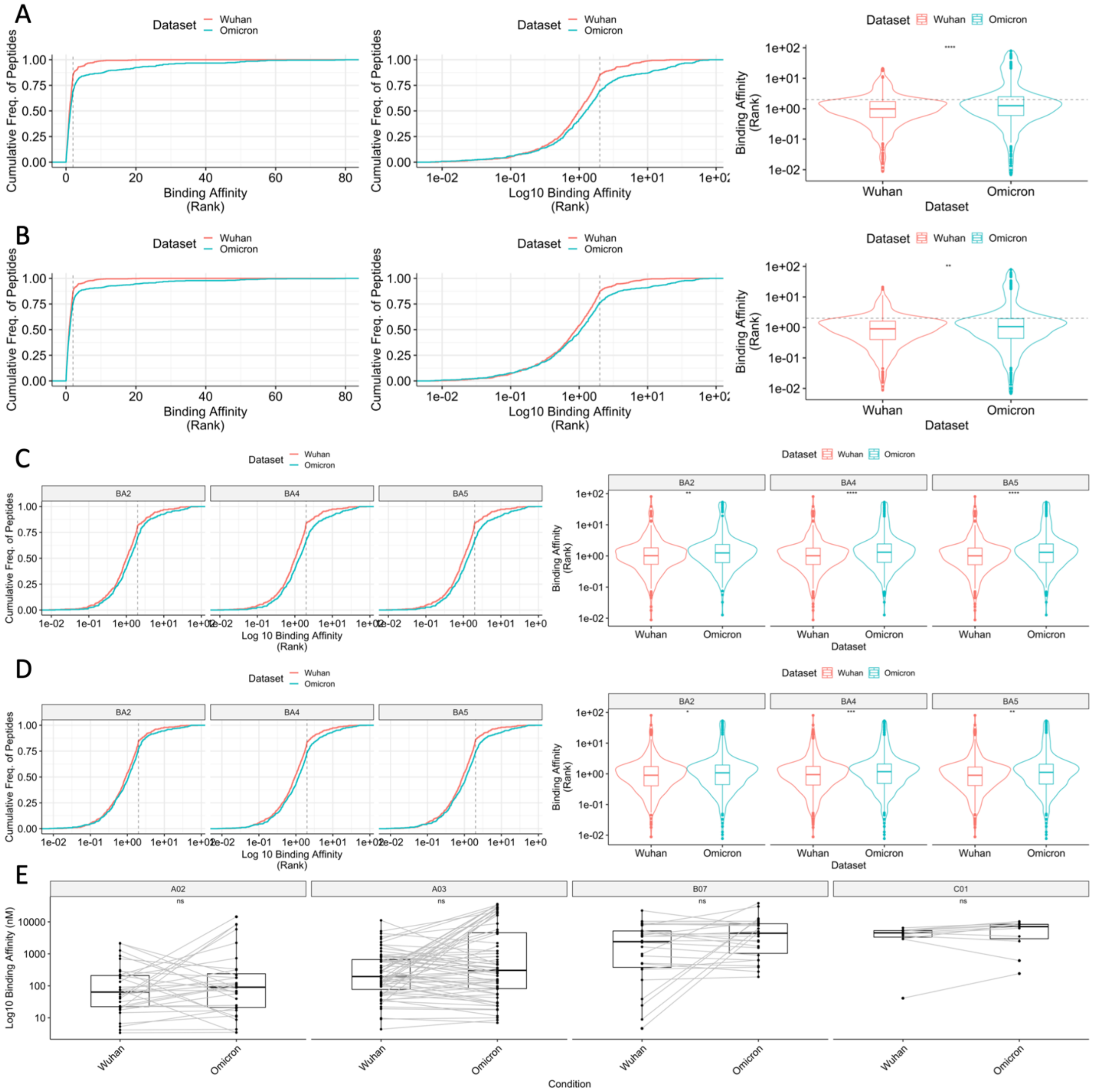

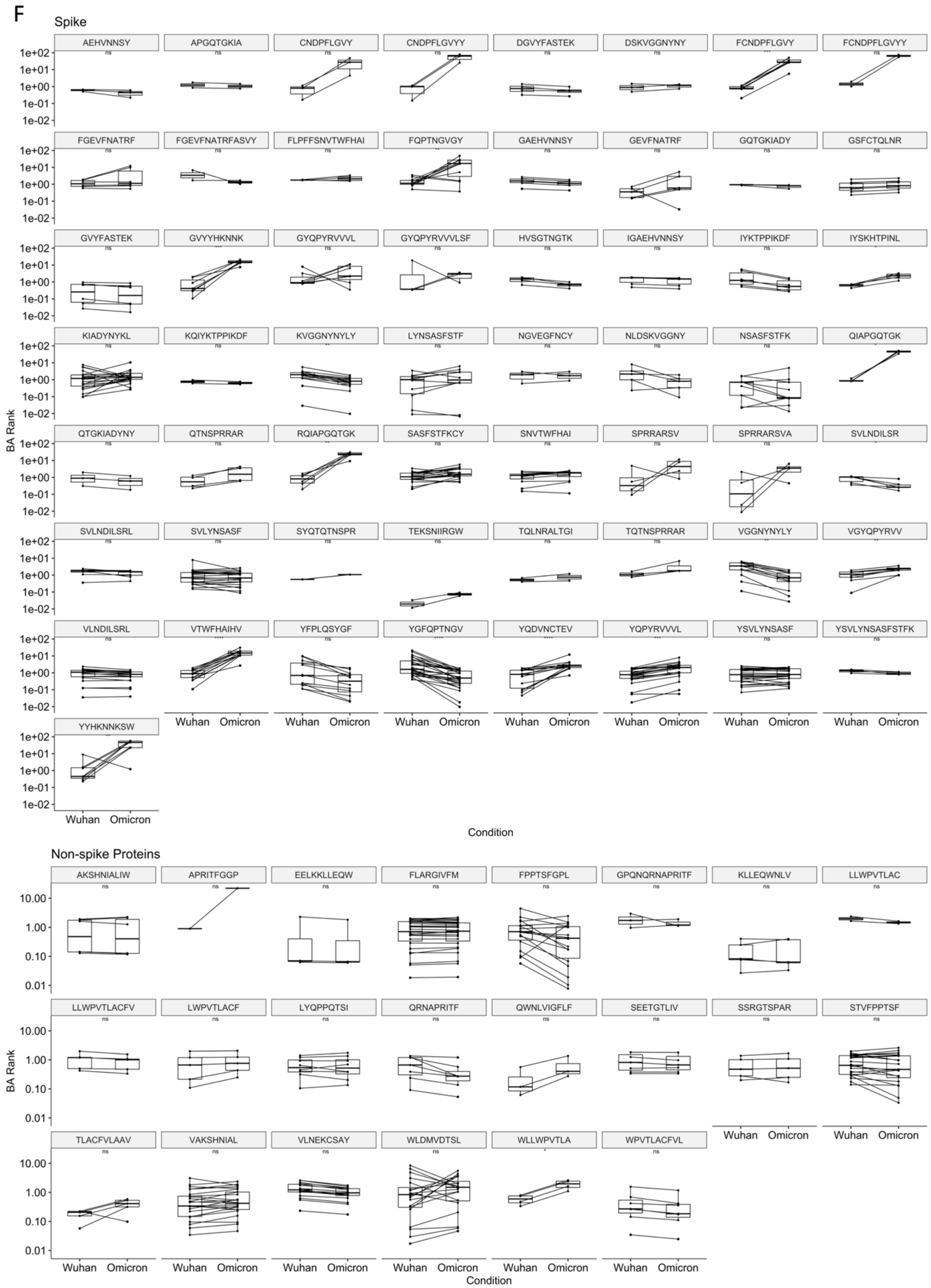


Supplementary Figure 2: A) ECDF, log10-ECDF, violin plots comparing netMHCpan binding affinity rank scores of spike-derived Wuhan vs. BA1 Omicron pMHC. Dashed line shows the binding rank threshold of 2.0 B) ECDF, log10 ECDF, violin plots comparing netMHCpan binding affinity rank scores of Wuhan vs. BA1 Omicron pMHC from any proteins exhibiting mutation in CD8+ T cell targets. Significance was assessed using a Wilcoxon rank test. Dashed line shows the binding rank threshold of 2.0 C) log10-ECDF and violin plots comparing netMHCpan binding affinity rank scores of spike-derived Wuhan vs. BA2, BA4, BA5 Omicron pMHC. Significance was assessed using a Wilcoxon rank test. Dashed line shows the binding rank threshold of 2.0 D) log10-ECDF and violin plots comparing netMHCpan binding affinity rank scores of Wuhan vs. BA2, BA4, BA5 Omicron pMHC from all affected proteins. Significance was assessed using a Wilcoxon rank test. Dashed line shows the binding rank threshold of 2.0 E) Paired boxplots for different HLA supertypes (HLA-A02, HLA-A03, HLA-B07, HLA-C01), comparing the binding affinity (rank score, log 10)

between Wuhan and BA1 Omicron. Significance was assessed using a paired Wilcoxon rank test. F) Paired boxplots for each pMHC, comparing the binding affinity rank score between Wuhan and BA1 Omicron. Significance was assessed using a paired Wilcoxon rank test.

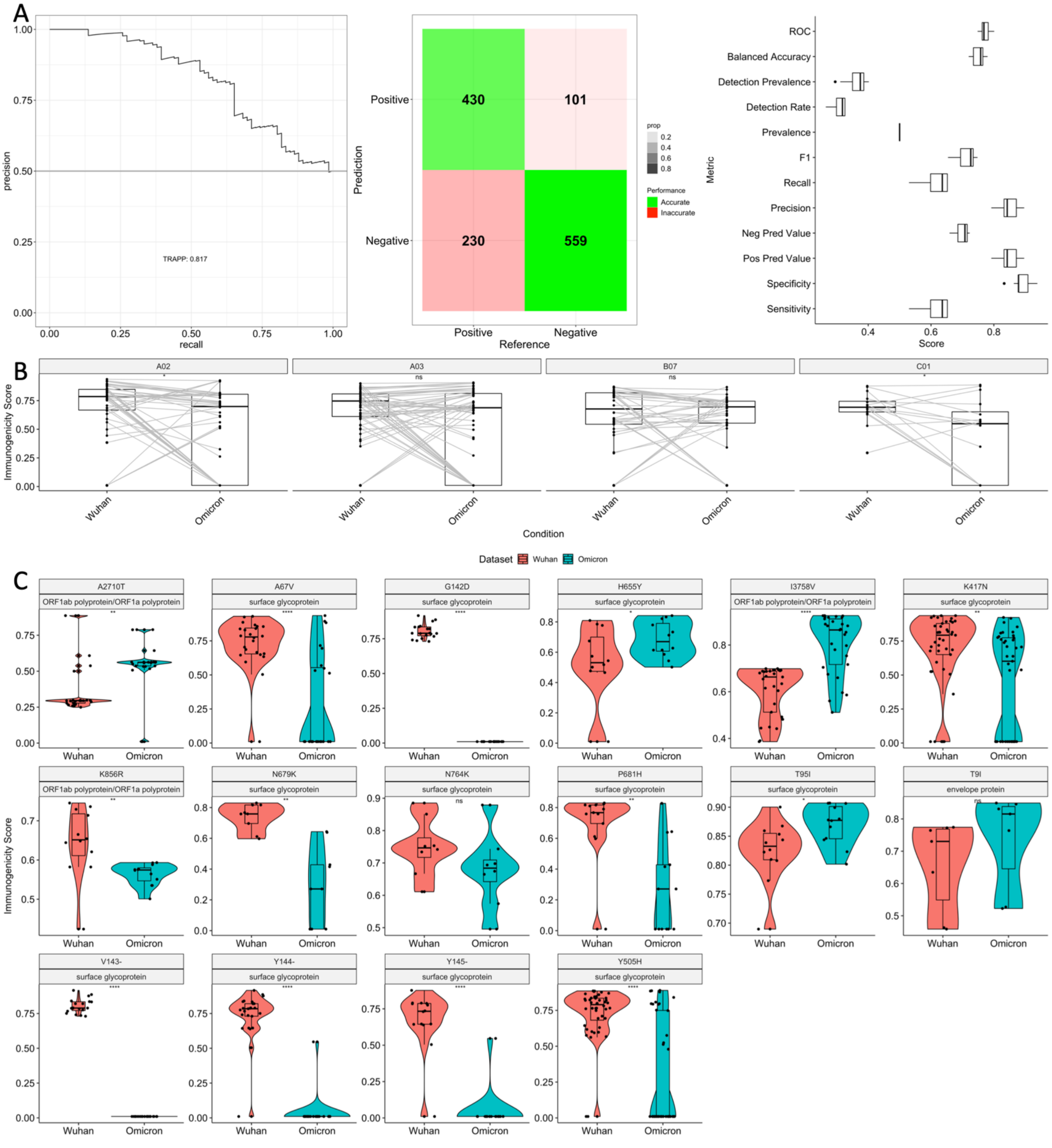

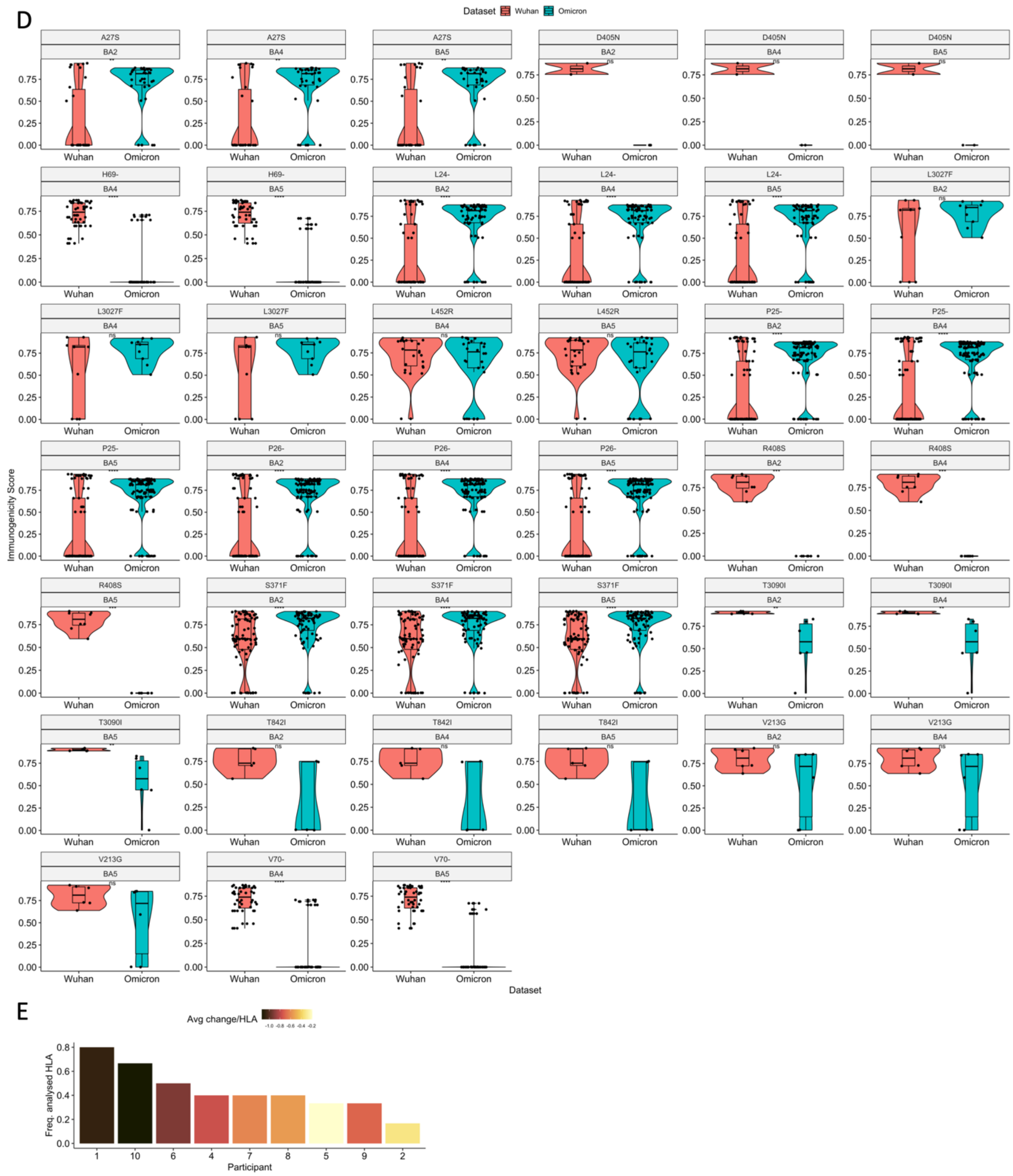


Supplementary Figure 3: A) Plots depicting the performance of TRAP against 66 known SARS-CoV-2 Wuhan Hu-1 CD8+ T cell targets with a mutation of Omicron. Negative scores were randomly sampled from a 10-fold cross-validation on the model training data. This means that each repetition (1..10), we randomly sampled 66 non-immunogenic epitopes and each time their TRAP scores are compared against the 66 Wuhan-mutated (positive) set. Precision recall plot (left) summarising these total data (n=660 pve, n=660 nve). Confusion matrix (middle) across each ten samplings. Boxplot showing the distributions of scores generated for multiple performance metrics. Each data point represents a metric generated from each sampling. B) Paired boxplots for different HLA supertypes (HLA-A02, HLA-A03, HLA-B07, HLA-C01), comparing the immunogenicity scores between Wuhan-mutated and BA1 Omicron epitopes. Significance was assessed using a Wilcoxon rank test. C) Jittered violin plots comparing the distribution of immunogenicity scores for pMHC affected by each BA1 mutation (those not selected for main text). Only mutations exhibiting significant differences are shown. Significance was assessed using Wilcoxon rank tests. D) Jittered violin plots comparing the distribution of immunogenicity scores for pMHC affected by each mutation unique to BA2, BA4 or BA5. Only mutations exhibiting significant differences are shown. Significance was assessed using a Wilcoxon rank test. E) Plot showing for Naranbhai et al's 10 patients whose T cell responses were impaired with Omicron, the frequency of their HLA genotype that bind MHC which exhibit a mean reduction in T cell immunogenicity (average log ratio MT/WT < -0.8 per HLA). This score (average log ratio per HLA) only considers CD8+ T cell targets with a mutation in Omicron. -0.8 was chosen as a cutoff to conservatively represent a reduction in immunogenicity where the addition of standard error was distinct from zero from all HLA.

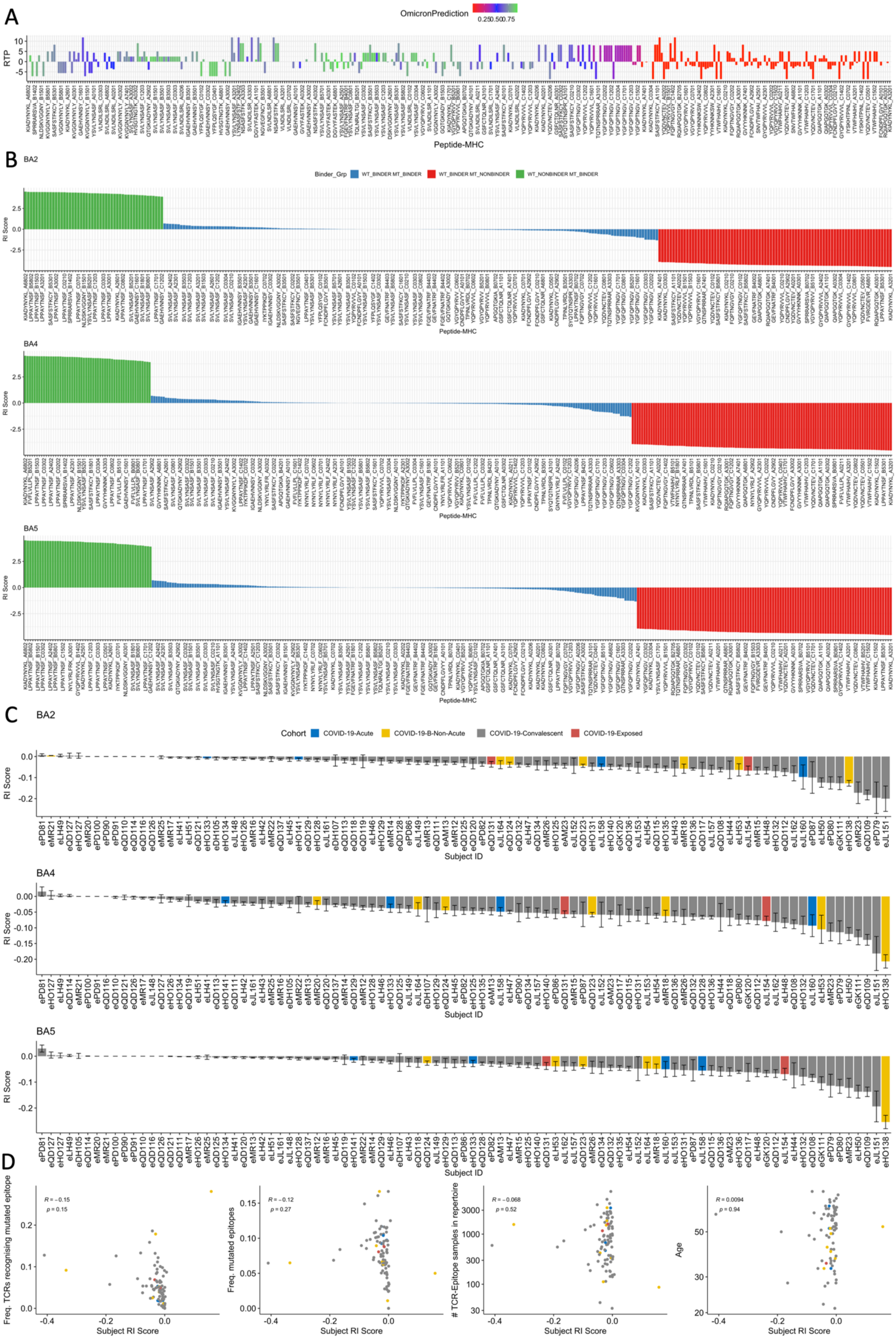
A
OmicronPrediction
RTP
Peptide-MHC
B
BA2
Binder_Grp
WT_BINDER MT_BINDER
WT_BINDER MT_NONBINDER
WT_NONBINDER MT_BINDER
RI Score
Peptide-MHC
BA4
RI Score
Peptide-MHC
BA5
RI Score
Peptide-MHC
C
BA2
Cohort
COVID-19-Acute
COVID-19-B-Non-Acute
COVID-19-Convalescent
COVID-19-Exposed
RI Score
Subject ID
BA4
RI Score
Subject ID
BA5
RI Score
Subject ID
D
R = −0.15
p = 0.15
Freq. TCRs recognising mutated epitope
Subject RI Score
R = −0.12
p = 0.27
Freq. mutated epitopes
Subject RI Score
R = −0.068
p = 0.52
### TCR-Epitope samples in repertoire
Subject RI Score
R = 0.0094
p = 0.94
Age
Subject RI Score

Supplementary Figure 4: A) Barplots showing for each mutated 9/10-mer BA1 CD8+ T cell target pMHC, the 'Relative TCR Promiscuity' score (RTP). RTP reflects the change in numbers of TCRs that bind each peptide. TITAN was used to generate binding predictions. Bars are colored coded by the immunogenicity prediction of the Omicron mutant. TITAN does not incorporate MHC, therefore RTP is produced for each peptide. Thus peptide 'AAA' bound to MHCs *k* and *l*, may have different RI scores but will have the same RTP. Plot is ordered by peptide-MHC RI score. B) Barplots showing the relative immunogenic potential (RI) score for each pMHC affected by a mutation in BA2 (top), BA4 (middle), BA5 (bottom) Omicron. C) Barplot showing the mean +/- standard error for 'pan-HLA RI' scores for each individual MIRA TCR-Epitope repertoire, for BA2, BA4, BA5. D) Scatter plots comparing BA1 MIRA subject average RI scores with i) frequency of their TCRs which recognise an epitope that is mutated in BA1 Omicron, ii) frequency of *unique* mutated epitopes recognised by TCRs in their repertoire, iii) the size, i.e., number of unique TCR-Epitope samples in their MIRA TCR repertoire, iv) their age.

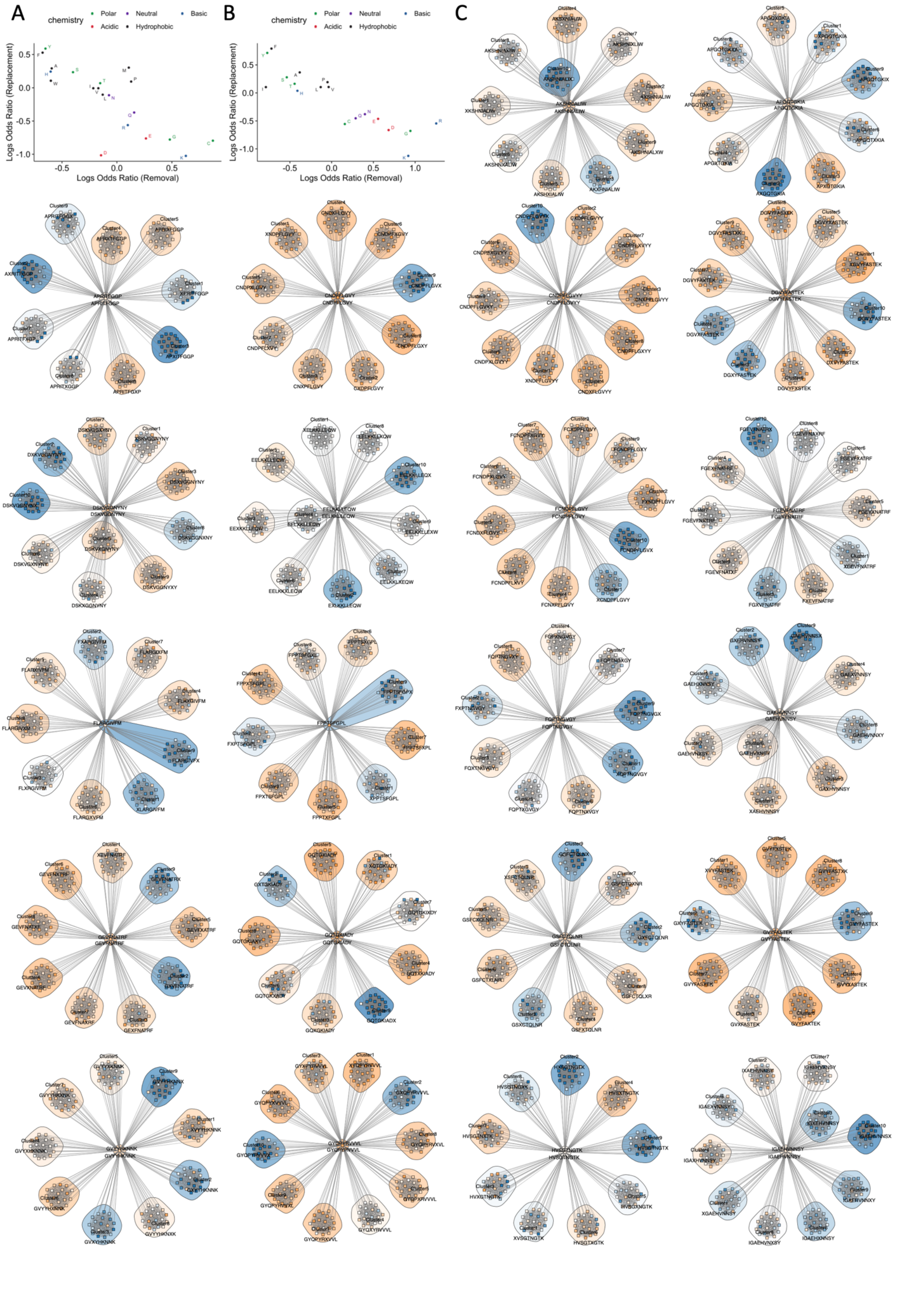
A
B
C
chemistry
Polar
Neutral
Basic
Acidic
Hydrophobic
Logs Odds Ratio (Replacement)
Logs Odds Ratio (Removal)

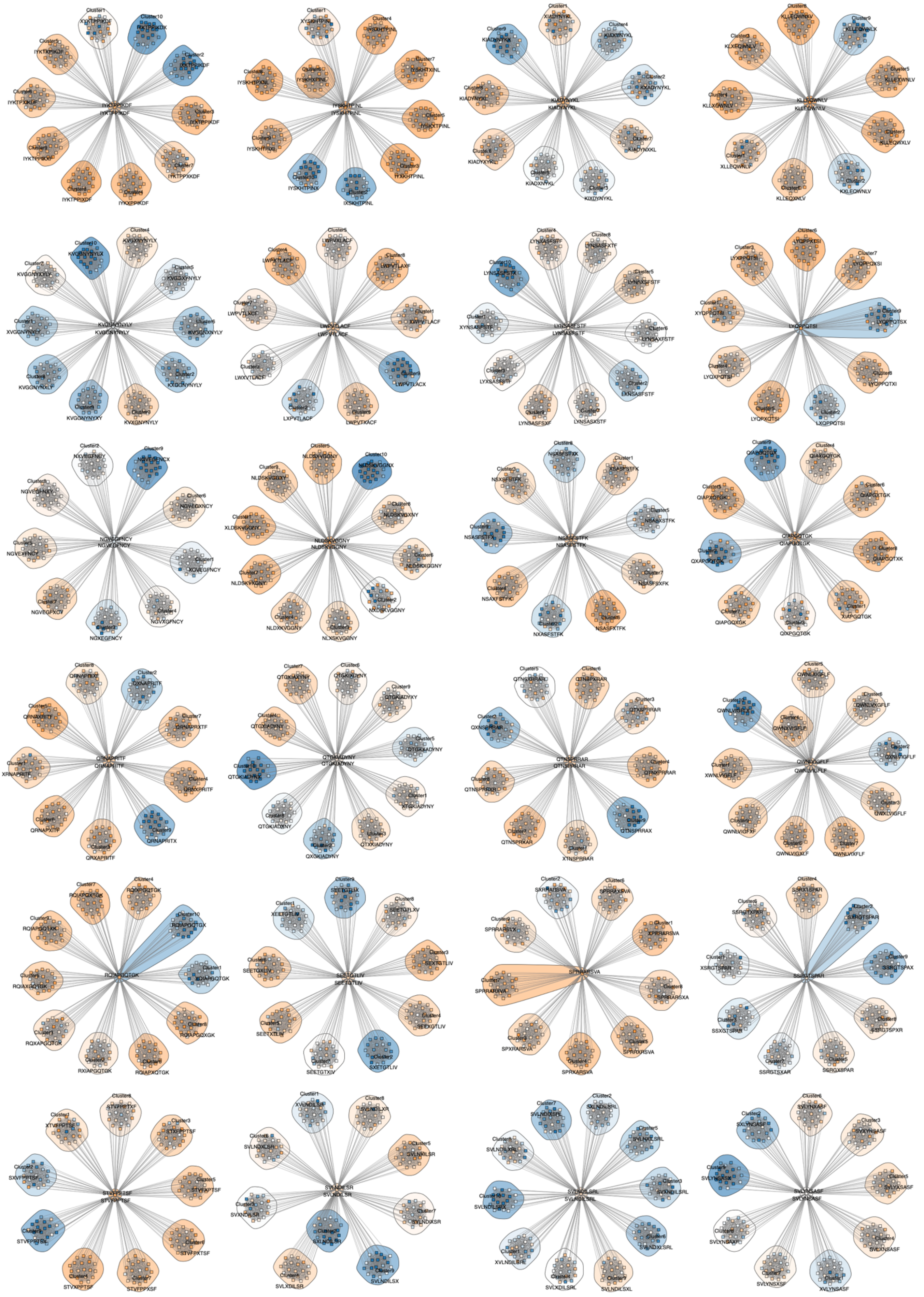
IYKTPPIKDF
IYSKHTPINL
KIADYNYKL
KLLEQWNLV
KVGGNYNYLY
LWPVTLACF
LYNSASFSTF
LYQPPQTSI
NGVEGFNCY
NLDSKVGGNY
NSASFSTFK
QIAPGQTGK
QRNAPRITF
QTGKIADYNY
QTNSPRRAR
QWNLVIGFLF
RQIAPGQTGK
SEETGTLIV
SPRRARSVA
SSRGTSPAR
STVFPPTSF
SVLNDILSR
SVLNDILSRL
SVLYNSASF

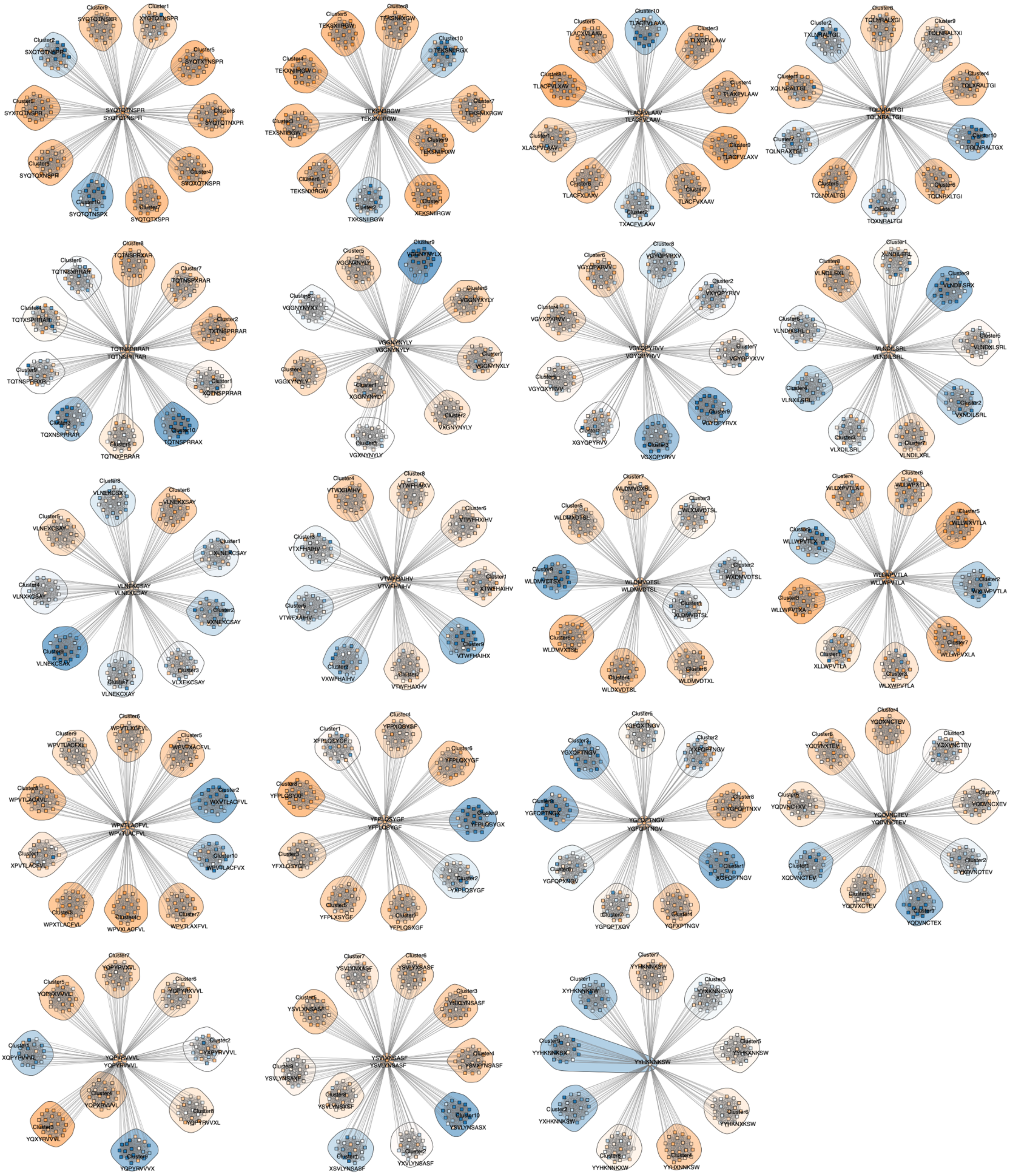

Supplementary Figure 5: In silico mutagenesis analysis A-B) Scatter plots showing the mean logs odds ratio changes in immunogenicity score (per residue) observed upon removing each amino acid from wildtype sequences (x-axis) vs. the mean logs odds ratio changes observed upon replacing (imputing) the amino acid into a mutant sequence for A) 9-mer contact positions, B) 10-mer contact positions. C) Neighbor network diagrams depicting mutational impact via trajectories from the wildtype peptide (center) to each single amino acid variant. Clusters and individual mutants are color labelled by immunogenicity score. Groups are clustered by location of the substitution. An X in the consensus sequence by the cluster indicates the position of the substitution that characterises the cluster.

#### Supplementary Methods

##### HLA alleles for antigen presentation prediction

HLA-A01:01,-A02:01,-A02:02,-A02:06,-A02:11,-A03:01,-A03:02,-A11:01,-A23:01,-A24:02,-A25:01,-A26:01,-A29:02,-A30:01,-A30:02,-A31:01,-A32:01,-A33:03, -A68:01, -A68:02, -A74:01,-B07:02,-B08:01,-B14:02,-B15:01,-B15:03,-B18:01,-B27:05,-B35:01,-B35:03,-B38:01,-B40:01,-B40:02,-B40:06,-B42:01,-B44:02,-B44:03,-B45:01,-B50:01,-B51:01,-B52:01,-B53:01,-B57:01,-B58:01,-B58:02,-C01:02,-C02:02,-C02:10,-C03:02,-C03:03,-C03:04,-C04:01,-C05:01,-C06:02,-C07:01,-C07:02,-C08:01,-C08:02,-C12:02,-C12:03,-C14:02,-C15:02,-C16:01,-C17:01

##### Identification of immunogenic Wuhan Hu-1 epitopes with a mutation in Omicron and its subvariants

Application of a filter that excluded any final matches that are too dissimilar from Wuhan Hu-1 was considered but not necessary. BA1 Omicron at the proteome and nucleotide level has one insertion that codes S:215EPE. At the proteome level, BA1 has two deletions: S:HV69del and S:VYY143del. We did not observe any epitopes affected by this insertion. For variant epitopes affected by deletions, we simply incorporated the next amino acid in the sequence e.g., [ABCDE~~F~~]G → [ABCDEG]. Thus, in this case, the deletion results in a single amino acid variant through a pairwise comparison. In all cases, to validate mutation identification, mutations we identified were compared with a list of defining mutations for each variant.

##### Re-training TITAN

We took the TITAN[1] pre-cursor which had been pre-trained on BindingDB, but not fine-tuned on TCR-epitope interaction data. For fine-tuning, functionally validated TCR data mapped to epitopes was gathered from the MIRA dataset, VDJdb and a recent publication from Pogorely et al. We excluded any samples containing epitopes of length>20 residues and unproductive TCRs. To generate full coding TCR sequences for fine-tuning, we employed the 'Thimble' function from the Stitchr package with default parameters for human analysis https://github.com/JamieHeather/stitchr. The script takes known V/J/CDR3 information, extracts the relevant nucleotide/coding sequence and generates a full coding sequence. 2320 TCRs were removed due to errors e.g., the TRBV/J gene annotation could not be matched with a database used for extracting the sequence. We thus curated full coding length sequences for 540,977 TCR samples. R package '*protr*' was used to confirm there were no TCR or epitope sequences containing non amino acid characters. SMILES descriptions of epitopes were generated using a custom script using the 'aas_to_smiles' function from the 'pytoda' Python package: https://paccmann.github.io/paccmann_datasets/api/pytoda.proteins.utils.html

Our total 'combined' data incorporated data from Pogorely et al., MIRA dataset, among others.

For training, we generated unique peptide and TCR IDs as required by TITAN. We labelled the functionally validated samples as '1', indicating their immunogenic/positive status. To avoid duplicate TCR-epitope observations, we extracted the unique subset of TCR-epitope samples, leaving 540,975 samples. We then extracted from our 'combined' dataset all remaining 4,577 samples from Pogorely et al and sampled 100,000 samples from the MIRA subset, after excluding any samples containing epitopes of length<9. We additionally extracted 22,666 samples from the original TITAN publication training data. At all stages, we excluded samples containing Wuhan epitopes with a mutation in BA1 Omicron (Wuhan-mutated). After excluding duplicates, our training data consisted of 118,707 distinct positive samples. For negative data, we generated every possible combination of TCR-epitopes from our positive data, excluded the positive samples, and then randomly sampled from the expanded set of data. This is a commonly used technique in this field, based on the assumption that it is unlikely to randomly select an epitope and TCR that bind and invoke a response.

TITAN was fine-tuned on these data using the full TCR sequence: V, J gene sequences and CDR3. TCRs were encoded using BLOSUM62. Due to superior performance in the authors' original study, we encoded the peptides using SMILES. Training occurred on 95% of the data while the remaining set was reserved for testing at each epoch. Training was performed using NVIDIA TITAN RTX gpu on a high-performance cluster.

The model was validated against 28,084 TCR-epitope samples (positive) consisting of 52 Wuhan-mutated epitopes. 28,084 negative samples were generated as described previously. The validation set TCRs may have been observed in the training data although the epitopes were entirely excluded. This is reasonable as we are interested in predicting binding of unseen epitopes – a more difficult problem - rather than unseen TCRs. Against these data, our fine-tuned model produced a ROC-AUC of 0.74 and a precision recall of 0.714, albeit with considerable variation in performance across epitopes.

Relative TCR Promiscuity Analysis

RTP is defined as the log ratio of # of TCRs that the mutant Omicron peptide binds vs. the wildtype from Wuhan. We therefore used the re-tuned TITAN model to predict binding between Wuhan-mutated peptides and their Omicron counterparts, against the same set of TCRs. Thus 69 9/10-mer Wuhan-mutated peptides and their 69 BA1 Omicron counterparts were compared against 162,390 TCRs from our combined dataset. Some of these TCRs were in the model training data, however Wuhan-mutated epitopes were entirely excluded. Again, this is reasonable as we are interested in the binding status of unseen epitopes rather than unseen TCR. To discriminate binding status vs. non-binding, we employed a conservative threshold of >0.85 for TITAN's output score (range 0-1). The number of TCRs that each assessed peptide was predicted to bind was counted. Any peptides not predicted to bind one single TCR (i.e a false negative), was imputed with a count of binding 1 TCR, as we know that these are immunogenic peptides thus there must be a single cognate T cell for this epitope.
